# All-in-one, Cas13d-based cell-specific gene knockdown system for zebrafish

**DOI:** 10.64898/2026.04.23.720501

**Authors:** Dongeun Heo, Cody L. Call, Jiakun Chen, Marc R. Freeman, Kelly R. Monk

## Abstract

Cell type-specific genetic manipulation is essential for dissecting neural circuits and glial biology *in vivo*; however, methods for achieving and validating cell-specific gene knockdown in zebrafish remain limited. Here, we introduce a CRISPR-Cas13d-based mRNA knockdown platform for zebrafish that leverages RNA targeting, rather than DNA editing, to enable efficient and flexible perturbation of gene function in defined cell types of interest. We engineered an all-in-one vector system for cell type-specific RfxCas13d expression and ubiquitous expression of crispr RNAs (crRNAs) and validated its utility across the major classes of CNS glia: astrocytes, oligodendrocytes, and microglia. Cas13d-mediated knockdown consistently achieved target mRNA depletion and produced heritable reproducible morphological and functional phenotypes, demonstrating the robustness of the approach. This approach complements permanent genome-editing methods such as CRISPR-Cas9 by providing a reliable, efficient, and scalable strategy for RNA-level perturbation. Our Cas13d toolkit expands the repertoire of zebrafish genetic technologies, offering a powerful resource for *in vivo* cell type-specific studies to uncover cell-autonomous mechanisms and possibility for cell type-specific genetic screens in development and disease.

## Main

Over the past few decades, the zebrafish (*Danio rerio*) has emerged as a powerful vertebrate model for uncovering the cellular and molecular mechanisms underlying nervous system development. External fertilization, rapid embryogenesis, and optical transparency of larval zebrafish allow direct visualization of developmental processes at high spatial and temporal resolution *in vivo*. Importantly, zebrafish represent the simplest vertebrate system that contains all major mammalian glial cell types, including protoplasmic astrocytes (Chen et al., 2020), oligodendrocyte lineage cells (Brösamle & Halpern, 2002; Kirby et al., 2006; Kucenas et al., 2008; Pogoda et al., 2006), and microglia (Herbomel et al., 1999; Xu et al., 2015). It is the only model organism that permits real-time monitoring of glial specification, migration, morphogenesis, and maturation in an intact animal; this provides a unique experimental advantage for understanding how complex glial cellular architecture and function emerge during development. In addition, the zebrafish genome exhibits substantial evolutionary conservation with humans. Approximately 70 percent of human genes have at least one zebrafish ortholog, including the majority of genes implicated in neurological and neurodevelopmental disorders (Howe et al., 2013). This conservation allows rigorous genetic interrogation of conserved molecular pathways that shape glial cell identity, maturation, and function that may be readily translationally relevant.

The foundation of zebrafish genetics was established through early work in the 1970s and 1980s (Streisinger et al., 1981; Walker & Streisinger, 1983), followed by large-scale forward mutagenesis screens in the 1990s (Driever et al., 1996; Grunwald & Streisinger, 1992; Haffter et al., 1996; Mullins et al., 1994). These N-ethyl-N-nitrosourea (ENU)-based screens generated thousands of random germline mutations and identified over one thousand reproducible developmental phenotypes affecting embryonic patterning, organogenesis, and nervous system development. Subsequent forward and modifier screens further expanded the zebrafish genetic toolkit, including insertional mutagenesis approaches and chemical genetic screens (Amsterdam et al., 2004; Peterson et al., 2000), which collectively enabled interrogation of vertebrate gene function across diverse biological processes. Together, these efforts firmly established zebrafish as a tractable and scalable vertebrate genetic system and provided fundamental insights into conserved developmental pathways. However, these early manipulations were constitutive and global, lacking spatial or temporal specificity, which limited their utility for studying gene function in defined neural cell populations.

Subsequent reverse genetic approaches, particularly morpholino-based gene knockdown (Nasevicius & Ekker, 2000), expanded experimental flexibility and facilitated hypothesis-driven studies using zebrafish. Despite their widespread use, morpholinos have been shown to frequently exhibit off-target effects, variable penetrance, and limited reproducibility (Joris et al., 2017; Kok et al., 2015; Stainier et al., 2017). Moreover, they did not provide stable or cell type-specific gene perturbation. Alternative strategies such as Targeting Induced Local Lesions in Genomes (TILLING) offered locus-directed mutagenesis but remained labor-intensive and inefficient (Wienholds et al., 2003).

The introduction of synthetic sequence-specific DNA nucleases (or programmable nucleases) marked a substantial advance in zebrafish genome engineering. Zinc finger nucleases (ZFNs) (Doyon et al., 2008; Meng et al., 2008; Urnov et al., 2005) and later transcription activator-like effector nucleases (TALENs) (Bedell et al., 2012; Miller et al., 2011; Sander et al., 2011) enabled target-specific DNA cleavage, improving the efficiency and precision of targeted mutagenesis. However, these approaches were technically complex, often time-consuming, and not readily scalable. The emergence of CRISPR-Cas9 technology (Jinek et al., 2012; Sternberg et al., 2014) transformed genetic accessibility in zebrafish by allowing highly efficient targeted mutagenesis through co-injection of Cas9 protein or mRNA and single-guide RNAs (sgRNAs) into single-cell zygotes (Chang et al., 2013; Hwang et al., 2013; Shah et al., 2015). This approach often produces detectable loss-of-function phenotypes in the injected (F_0_) “crispant” animals, enabling rapid functional interrogation. Despite these advances, DNA double-strand break-based genome editing introduces intrinsic biological and technical challenges. Global disruption of essential genes often leads to early embryonic lethality, precluding investigation of later roles in nervous system development and function.

Therefore, cell- and tissue-specific Cas9 strategies have been developed in zebrafish, including promoter-driven expression of Cas9 and Gal4/UAS-based binary systems (Ablain & Zon, 2016; De Santis et al., 2016; Di Donato et al., 2016; Li et al., 2024). These approaches significantly expanded the genetic toolbox for zebrafish neurobiology research and enabled conditional mutagenesis in defined cell populations. However, Cas9-mediated editing still relies on double-strand DNA breaks, which generate heterogeneous in/dels—including silent and missense mutations—that can drastically vary between targeted cells in the absence of a donor template, confounding quantitative analyses and obscuring subtle cell-autonomous phenotypes in neural tissue. In addition, double-strand breaks activate DNA damage responses, including p53-mediated apoptosis (Haapaniemi et al., 2018), and induce large chromosomal aberrations with the usage of multiple sgRNAs (Kosicki et al., 2018; Shin et al., 2017), which can confound phenotypic interpretation without appropriate *post hoc* validations. However, rigorous validation of gene loss remains challenging as Cas9-mediated gene targeting does not uniformly eliminate transcript production (Smits et al., 2019; Tuladhar et al., 2019), leads to mosaicism (Mehravar et al., 2019), and confirmation of protein depletion frequently relies on antibodies that are unavailable for many zebrafish targets. Furthermore, direct visualization of genetically perturbed cells with tissue-specific Cas9 remains technically challenging. Strategies that couple Cas9 expression to fluorescent reporters, such as Cas9-2A-EGFP bicistronic expression constructs (Ablain & Zon, 2016; De Santis et al., 2016; Di Donato et al., 2016), can yield weak or inconsistent labeling, likely due to the large size of the Cas9 protein.

Ideally, we need a complementary genetic strategy that achieves reliable, quantifiable, and cell type-specific gene perturbation while avoiding permanent genomic alteration. An ideal system would allow scalable targeting of multiple genes across diverse neural cell populations and would be compatible with stable transgenic lines. To this end, we developed an all-in-one vector platform based on CRISPR-Cas13d, an RNA-guided ribonuclease that targets single-stranded RNA transcripts rather than genomic DNA (Abudayyeh et al., 2017; Gupta et al., 2022; Kushawah et al., 2020; Powell et al., 2022). Cas13d mediates transcript depletion at the post-transcriptional level, thereby avoiding double-strand breaks and permanent DNA modification. The enzyme is compact and programmable with a single-base precision, and functions independently of the endogenous Dicer pathway, distinguishing it mechanistically from traditional RNA interference (RNAi) approaches (Shembrey et al., 2024; Wessels et al., 2020, 2024). Furthermore, Cas13d-mediated knockdown is reversible (Lv et al., 2022) and has been successfully adapted across multiple model systems, including zebrafish (Kushawah et al., 2020; Moreno-Sánchez et al., 2025), where it supports efficient *in vivo* gene knockdown with minimal toxicity.

By combining Cas13d with the optimized Gal4-activator (KalTA4)/UAS binary system (Distel et al., 2009) and a multiplex gRNA expression system (Yin et al., 2015), we developed a cell type-specific Cas13d platform that enables flexible targeted gene knockdown in defined glial populations within the nervous system. In addition, we implemented a complementary approach in which Cas13d expression is driven directly by a cell type-specific promoter, thereby eliminating the need for an established KalTA4 transgenic zebrafish line.

Notably, this system enables robust *in vivo* visualization of Cas13d-targeted cells, providing a powerful genetic tool for tracking and quantifying gene perturbation at single-cell resolution. We demonstrate that this approach produces consistent RNA transcript depletion, reproducible cellular phenotypes in different cell types, and stable inheritance in transgenic lines. Collectively, this RNA-targeting genetic toolkit expands the experimental repertoire available for vertebrate neurobiology and provides a flexible and scalable framework for interrogating genetically redundant pathways, dissecting glial cell biology, and conducting cell type-specific functional genetic screens *in vivo*.

## Results

### Adaptation of a global Cas13d strategy for an all-in-one, cell type-specific system

CRISPR-Cas13d has been successfully adapted for zebrafish to achieve transcript-level knockdown of genes of interest during early development (Kushawah et al., 2020; Moreno-Sánchez et al., 2025). Building on this foundation, we developed an all-in-one construct for KalTA4-UAS-based, cell type-specific expression (**Fig. 1**). To do so, we cloned *UAS-RfxCas13d* into the *pDest-Tol2-4sgRNA-Cas9* Gateway destination vector (**Fig. 1a**), a backbone originally designed for Cas9-based gene manipulations in zebrafish (Yin et al., 2015). This destination vector provides several features that are advantageous for efficient cloning process: (1) *BsaI* sites that enable rapid one-pot, Golden Gate cloning; (2) layers of negative and positive selection, including *CmR* and *ccdA/B* cassettes, which greatly reduce non-specific products; and (3) Tol2 inverted terminal repeats (ITRs), allowing for Tol2 transposase-mediated genomic integration. Because plasmid dilution during early zebrafish embryogenesis poses a challenge for stable and cell-specific expression in glia, which are generated later in development, Tol2-mediated transgenesis is essential to achieve robust and heritable gene expression. The relatively small size of Cas13d (930 amino acids) compared with Cas9 (1,400 amino acids) also keeps the final construct within the cargo capacity well tolerated by the Tol2 system.

**Fig. 1.**
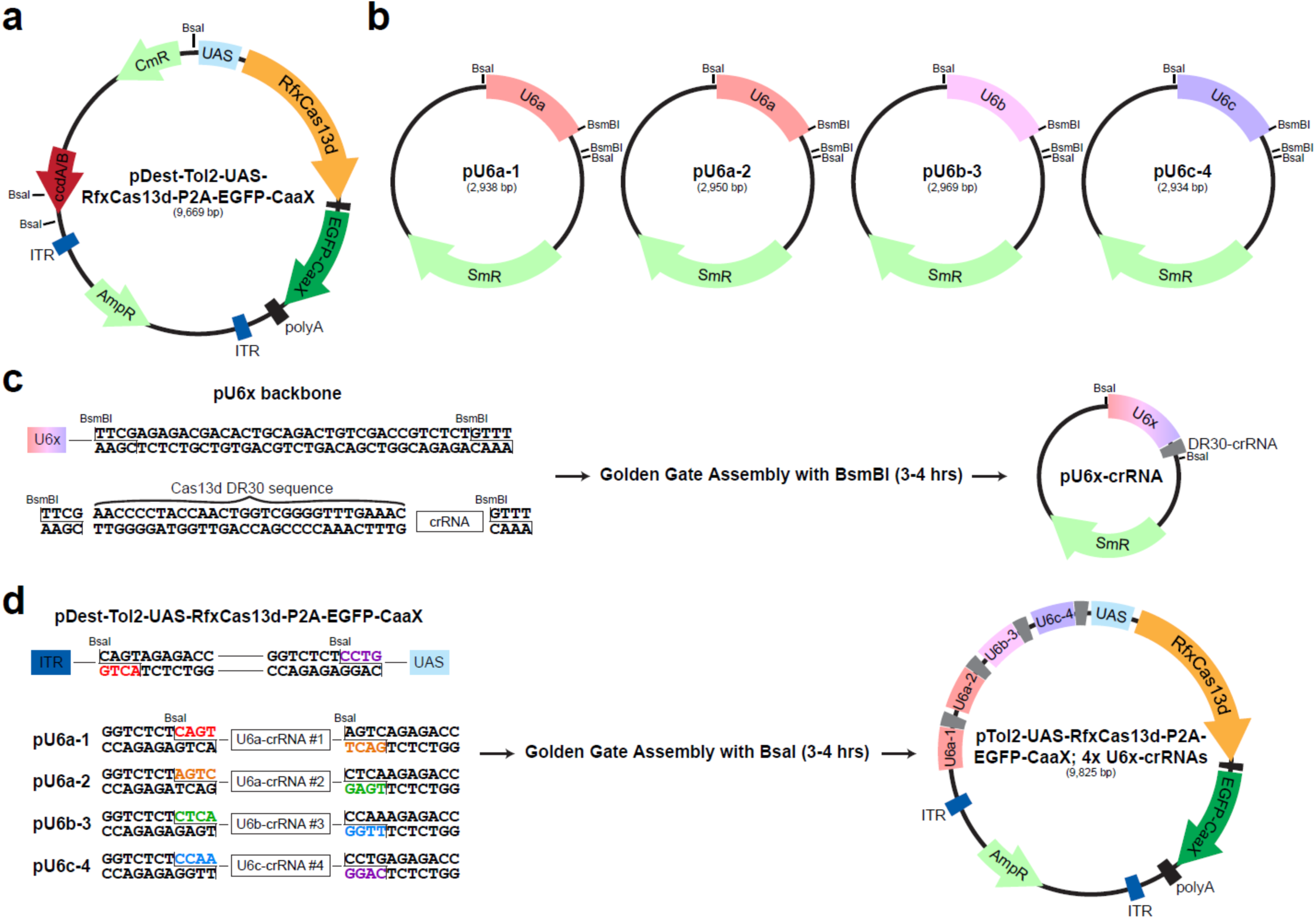
Development of all-in-one, cell type-specific Cas13d system. **a**. A schematic diagram of the plasmid map of destination vector: *pDest-Tol2-UAS-RfxCas13d-P2A-EGFP-CaaX*. RfxCas13d (orange) and EGFP-CaaX (dark green) is expressed under the control of UAS element. *BsaI* sites flank the ccdA/B and CmR (Chloramphenicol resistance gene; light green) sequences. **b**. Schematics of the four pU6x plasmid maps. Three distinct zebrafish U6 promoters (*U6a*, *U6b*, and *U6c*) were used to drive the expression of four different target crRNAs. The plasmids contain two *BsmBI* sites for the incorporation of DR30-crRNA and two *BsaI* sites for integration into the destination vector. **c**. Annealed pair of primers that contain *BsmBI* cut sites, DR30 sequence, and target-specific crRNA sequence, are cloned into the pU6x plasmids using the one-pot, Golden Gate Assembly protocol. **d**. Four pU6x-crRNA plasmids are cloned into the destination vector using the Golden Gate Assembly with the *BsaI* restriction enzyme. The schematic of a completed cell type-specific Cas13d plasmid is shown on the right.

To enable visualization of targeted cells *in vivo*, we incorporated a membrane-targeted EGFP reporter. Specifically, we fused an *EGFP-CaaX* sequence downstream of *RfxCas13d* via a P2A “self-cleaving” peptide (**Fig. 1a**). This bicistronic strategy allows both the tracking of targeted cells and visualization of cellular morphology. We next adapted the existing *pU6x* vectors, originally developed for multiplex expression of Cas9 sgRNAs (Yin et al., 2015), for Cas13d to enable simultaneous expression of multiple crRNAs and enhance knockdown efficiency. To accommodate crRNAs for Cas13d, we removed the *tracrRNA* scaffold sequences at the 3′ end while preserving *BsmBI* cloning sites for insertion of *DR30-crRNA* sequences and *BsaI* sites compatible with the Gateway destination vector (**Fig. 1b**). For crRNA design, pairs of primers containing *BsmBI* recognition sequences, the DR30 direct repeat, and a 22-23 bp targeting sequence of interest were synthesized and annealed (**Fig. 1c**). Target sequences were selected using RNAfold software-based *in silico* secondary structure predictions (Gruber et al., 2008), following the previously established global Cas13d zebrafish protocol (Kushawah et al., 2020; Moreno-Sánchez et al., 2025). Because Cas13d preferentially cleaves single-stranded regions of RNA, we designed crRNAs against regions predicted to have high single-stranded probability to maximize efficiency.

One-pot, Golden Gate cloning enabled efficient insertion of candidate DR30-crRNA sequences into *pU6x* vectors within 3-4 hours, with a typical success rate exceeding 90%. After constructing individual crRNA plasmids, we assembled them into the destination vector using a second Golden Gate reaction with *BsaI*. This step allows the simultaneous directional insertion of four *pU6x-crRNA* cassettes into a single *pTol2-UAS-Cas13d* expression construct (**Fig. 1d**). The ccdA/B negative selection system ensured exceptionally high cloning fidelity, with nearly all recovered colonies carrying the correct insert (∼100%). This modular design also permits rapid generation of different combinations of crRNAs, enabling scalable knockdown of multiple genes or targeting of distinct regions of the same transcript. Together, this optimized cloning workflow (**Extended Data Fig. 1**) streamlines the generation of cell type-specific Cas13d constructs for zebrafish, ensuring efficient transgenesis, robust visualization of targeted cells, and flexible multiplexing of crRNAs.

### Functional validation of cell type-specific Cas13d system in astrocytes

To evaluate the efficacy of our cell type-specific Cas13d system, we focused on *s1pr1*, a gene previously shown to regulate astrocyte morphology and growth in larval zebrafish (Chen et al., 2024). Global loss-of-function mutations in *s1pr1* result in a significantly reduced astrocyte cellular volume. Given the enriched expression of *s1pr1* in astroglia across both zebrafish and mammals (Hrvatin et al., 2018; Zhang et al., 2014), it is hypothesized that it acts cell-autonomously in astrocytes. However, reliable cell-specific perturbation has been difficult to achieve with existing techniques, motivating us to test whether our Cas13d system could recapitulate astrocyte morphological phenotypes observed in global *s1pr1* mutants in a targeted manner. To this end, we designed four independent crRNAs against the protein-coding region of *s1pr1* mRNA (**Fig. 2a**). To enable within-animal comparisons, we also employed the Tg(*UAS:myr-mScarlet*) reporter line (Li et al., 2024) (**Fig. 2b**) where we can image both the control and targeted cells in the same field of view within the same experimental animal. As an initial validation, we asked whether our cell type-specific Cas13d system could reduce the abundance of *s1pr1* mRNA transcripts. A longstanding limitation of Cas9-based approaches has been the difficulty of validating loss of function, as many Cas9-mediated genetic mutations do not reliably trigger nonsense-mediated decay (Smits et al., 2019; Tuladhar et al., 2019), and antibodies suitable for detection in zebrafish are often unavailable. By contrast, the RNA-targeting activity of Cas13d allows direct assessment at the transcript level in targeted cells using whole-mount fluorescence *in situ* hybridization (FISH) (Choi et al., 2018; Ibarra-García-Padilla et al., 2021). Using the Tg(*slc1a3b:KalTA4*) driver line (Lambert et al., 2025) to selectively target astrocytes in the ventral spinal cord, we observed a significant reduction (∼50%) in *s1pr1* mRNA within targeted astrocytes at 5 days post-fertilization (dpf) (**Fig. 2c,d**). Although depletion was not complete—likely due to continued transcription of new *s1pr1* mRNA transcripts—the knockdown was robust and statistically significant.

**Fig. 2.**
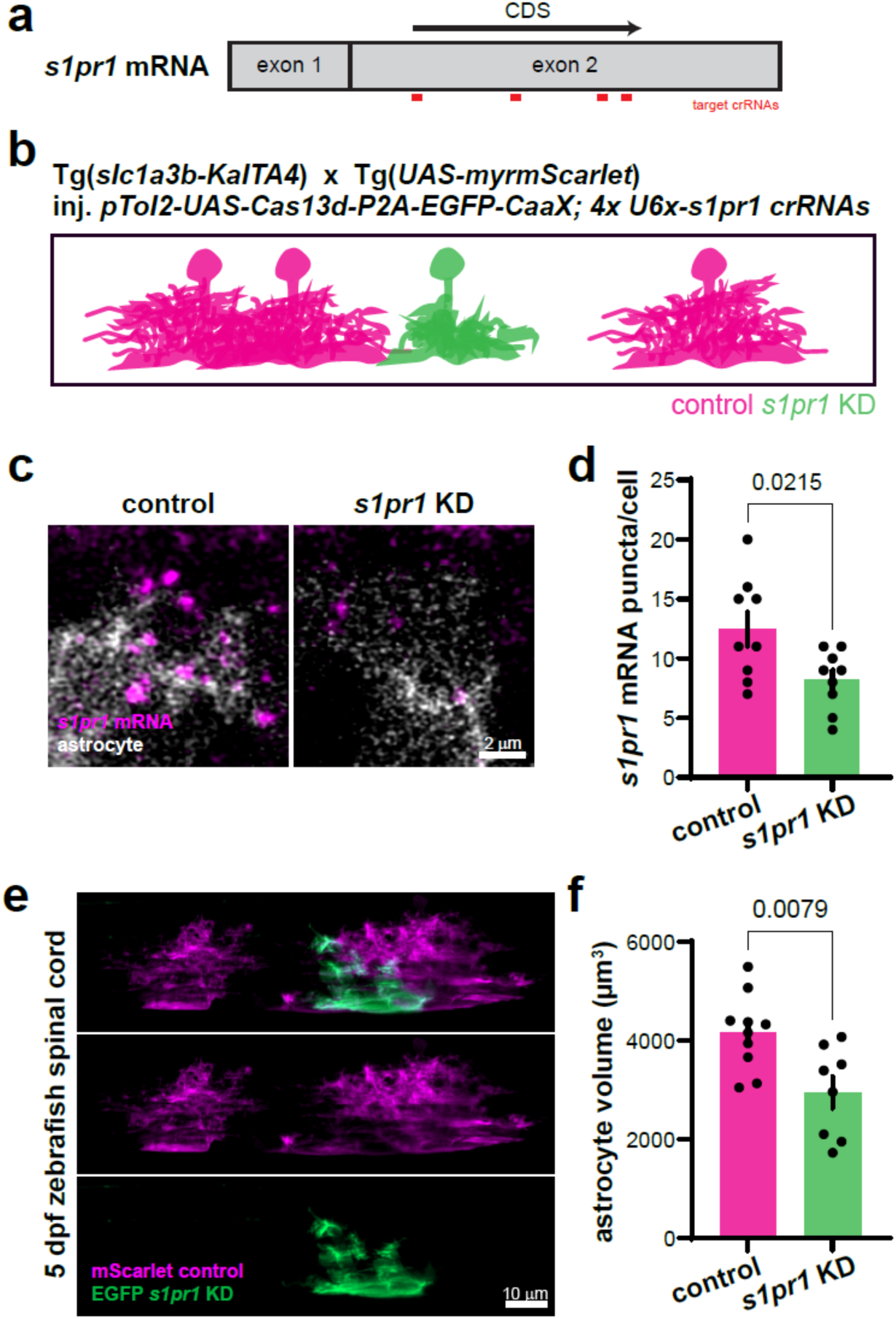
Functional validation of the cell-specific Cas13d system in astrocytes. **a**. Four distinct crRNAs (in red) were chosen against the protein-coding sequence (CDS) of the mature mRNA sequence of zebrafish *s1pr1*. **b**. Schematic of experimental design. Single-cell embryos of Tg(*slc1a3b-KalTA4*) crossed to Tg(*UAS-myrmScarlet*) were injected with *pTol2-UAS-Cas13d-P2A-EGFP-CaaX; 4x U6x-s1pr1 crRNAs*, which allows visualization and quantification of mScarlet+ control astrocytes (magenta) and EGFP+ targeted astrocytes (green) within the same animal. **c**. Representative whole-mount fluorescence *in situ* hybridization (FISH) images of control and *s1pr1* KD astrocyte soma with *s1pr1* mRNA puncta. **d**. Quantification of *s1pr1* mRNA puncta per astrocyte cell body. N=9 astrocytes/genotype. **e**. Representative confocal microscope images of EGFP+, targeted astrocytes (green) in the same field of view as mScarlet+, control astrocytes (magenta). **f**. Quantification of astrocyte volume determined by Imaris 3D Surface rendering. N=9 astrocytes/control, N=8 astrocytes/*s1pr1* KD.

We next tested whether Cas13d-mediated *s1pr1* knockdown could recapitulate the morphological phenotype observed in global mutants. Consistent with our expectations, the cellular volume of targeted ventral astrocytes was significantly reduced upon *s1pr1* knockdown using the cell type-specific Cas13d system compared to that of the controls (**Fig. 2e,f**). The degree of reduction in volume (∼30%) is comparable to previously reported volume change with the global *s1pr1* loss-of-function mutant (∼40%) (Chen et al., 2024). This demonstrates that Cas13d can efficiently knockdown *s1pr1* in targeted astrocytes and functionally reproduce morphological phenotypes previously associated with the global loss-of-function approaches. Finally, to assess the stability and heritability of this approach, we generated multiple stable transgenic lines, including Tg(*UAS:Cas13d-P2A-EGFP-CaaX; 4xU6x-s1pr1 crRNAs*), Tg(*UAS:Cas13d-P2A-EGFP-CaaX; 4xU6x-tracrRNAs*), and Tg(*UAS:Cas13d-P2A-EGFP-CaaX; 4xU6x-dsRed crRNAs*). Analysis of the F_2_ progeny using Tg(*slc1a3b-KalTA4*) driver line confirmed the presence of the astrocyte morphological phenotype (**Extended Data Fig. 2a,b**). To address prior reports of UAS transgene silencing in zebrafish, we performed whole-mount FISH (Choi et al., 2018; Ibarra-García-Padilla et al., 2021) for *egfp* mRNA and found that a subset of EGFP+ ventral astrocytes in *s1pr1* knockdown (KD) animals (Tg(*slc1a3b-KalTA4, UAS-Cas13d-P2A-EGFP-CaaX; U6x-s1pr1 crRNAs*)) lacked detectable *egfp* mRNA signal (**Extended Data Fig. 2c**). This suggests that, in a fraction of cells, transcription of the UAS-driven transgene (*Cas13d-P2A-EGFP-CaaX*) is attenuated due to silencing despite detectable EGFP fluorescence. Notably, this effect was infrequent, occurring in approximately 10% of EGFP+ cells (5 of 48 cells), indicating that UAS silencing represents a minor but measurable source of variability when using KalTA4-UAS binary expression system.

Together, these results establish that our all-in-one, cell type-specific Cas13d system enables efficient, quantifiable, and heritable knockdown in zebrafish astrocytes. This approach thus provides a powerful tool for interrogating glial gene function with spatial precision and across generations.

### Cell type-specific Cas13d system in oligodendrocyte lineage cells

Larval zebrafish provide a uniquely powerful system for investigating oligodendrocyte lineage progression, as oligodendrocyte precursor cells (OPCs) can be visualized differentiating into mature, myelinating oligodendrocytes in real time *in vivo* (Brösamle & Halpern, 2002; Czopka et al., 2013; Kirby et al., 2006; Kucenas et al., 2008; Pogoda et al., 2006). In the larval spinal cord, oligodendrogenesis is highly efficient: the vast majority of OPCs that initiate differentiation successfully mature and survive (Almeida & Lyons, 2016). This near-complete efficiency establishes a stringent baseline against which subtle perturbations in differentiation or survival can be detected with high sensitivity.

We therefore used this system to test whether our cell type-specific Cas13d platform could functionally knock down a gene with a well-established role in oligodendrocyte biology. TDP-43, encoded by *Tardbp*, is a ubiquitously expressed DNA/RNA-binding protein that regulates RNA processing and transcriptional integrity (Hanson et al., 2011). Conditional deletion studies in mice have demonstrated that TDP-43 is required for OPC survival and long-term oligodendrocyte maintenance (Heo et al., 2022; Wang et al., 2018). However, owing to limitations in lineage-restricted genetic drivers, it remains unresolved whether TDP-43 directly controls the differentiation capacity of OPCs *in vivo*. To address this question, we performed oligodendrocyte lineage-specific knockdown of *tardbp* in zebrafish. The zebrafish genome contains two orthologs, *tardbpa* (formerly *tardbpl*) and *tardbpb* (formerly *tardbp*). Combined loss of both paralogs results in developmental lethality, whereas individual loss leads to reduced adult survival, consistent with partial redundancy and genetic compensation (Schmid et al., 2013). To overcome this redundancy, we designed two crRNAs targeting the protein-coding region of *tardbpa* and two targeting *tardbpb*, for a total of four independent crRNAs, leveraging the all-in-one Cas13d system to enable simultaneous multiplexed targeting, thereby allowing efficient knockdown of both paralogs within a single construct (**Fig. 3a**).

**Fig. 3.**
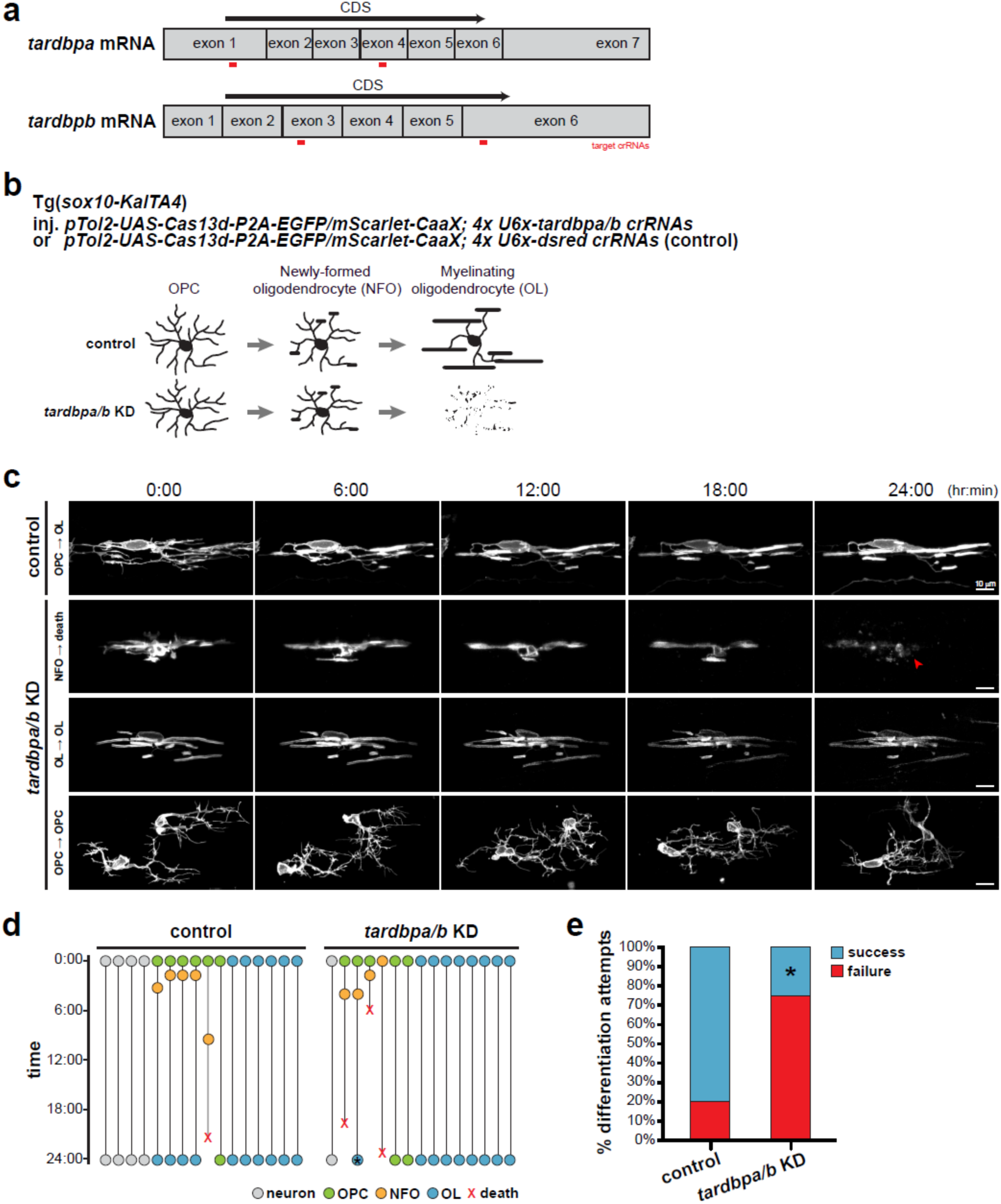
Functional validation of the cell-specific Cas13d system in oligodendrocyte lineage cells. **a**. Four distinct crRNAs (in red) were designed to target the protein-coding sequence (CDS) of the mature mRNA sequence of zebrafish *tardbpa* (2x) and *tardbpb* (2x). **b**. Schematic of experimental design. Single-cell embryos from Tg(*sox10-KalTA4*) were injected with *pTol2-UAS-Cas13d-P2A-EGFP/mScarlet-CaaX; 4x U6x-tardbpa/b crRNAs* or *pTol2-UAS-Cas13d-P2A-EGFP/mScarlet-CaaX; 4x U6x-dsred crRNAs* (control). **c**. Representative images of targeted oligodendrocyte lineage cells at 3 dpf, imaged over 24 hours at 2-hour intervals. The red arrow indicates a degenerating newly-formed oligodendrocyte (NFO) following *tardbpa/b* KD. **d**. Plot of individual cell fates in control and *tardbpa/b* KD conditions. All targeted neurons (gray) and myelinating oligodendrocytes (OLs; blue) survive throughout the imaging period. In contrast, differentiating OPCs and NFOs (orange) underwent degeneration (red X) following *tardbpa/b* KD. The asterisk (*) indicate an aberrant oligodendrocyte morphology. **e**. Quantification of OPC differentiation outcomes, categorized as successful (blue) or failed (red). N=5 NFOs/control, N=4 NFOs/*tardbpa/b* KD.

Using the Tg(*sox10-KalTA4*) driver line to restrict Cas13d expression to oligodendrocyte lineage cells and some interneurons (Almeida & Lyons, 2015) (**Fig. 3b**, **Extended Data Fig. 3**), we assessed the stage-specific consequences of TDP-43 depletion by performing longitudinal imaging at 3 dpf. We found that neurons, OPCs, and mature myelinating oligodendrocytes are largely resistant to *tardbpa/b* knockdown (**Fig. 3c, Extended Data Fig. 3**), indicating that the Cas13d construct overexpression alone does not induce widespread, non-specific toxicity. Strikingly, differentiating OPCs and newly formed oligodendrocytes displayed selective vulnerability upon TDP-43 depletion, undergoing cell death during the cellular transition to myelinating oligodendrocytes (**Fig. 3c,d**). These findings provide *in vivo* evidence that TDP-43 directly controls oligodendrocyte lineage progression in a cell-autonomous manner. In addition, complementary work from our lab has applied the Cas13d system to knock down *piezo1* and *piezo2* specifically within oligodendrocyte lineage cells to define the role of PIEZO channels in oligodendrocyte maturation and myelination (Coombs et al., 2026). Together, these results establish the utility of our cell type-specific Cas13d system for defining stage-specific gene function within the oligodendrocyte lineage *in vivo* and provide direct evidence that TDP-43 is required for OPC differentiation and the survival of newly-formed oligodendrocytes.

### Cell type-specific Cas13d system in microglia

Microglia are the resident immune cells of the CNS and play critical roles in neurodevelopment, synaptic remodeling, and debris clearance (Michell-Robinson et al., 2015; Paolicelli et al., 2011; Schafer et al., 2012). Increasing evidence also implicates microglia as central contributors to neurodegenerative disease pathogenesis (Hansen et al., 2018; Prinz et al., 2019). Despite their importance, rapid cell type-specific genetic manipulation of microglia *in vivo* remains challenging in mammalian systems, as currently available viral approaches are limited by low transduction efficiency or poor specificity (Balcaitis et al., 2005; Cucchiarini et al., 2003; Maes et al., 2019). Accordingly, there remains an enormous need for an efficient and scalable strategy to interrogate gene function in microglia. We therefore evaluated whether our all-in-one, cell type-specific Cas13d system could address this gap.

To adapt our platform for microglial targeting, we drove Cas13d expression under the control of the microglia/macrophage-specific promoter *mpeg1* (Ellett et al., 2011a). In contrast to the KalTA4/UAS binary system used in previous experiments for astrocytes and oligodendrocyte lineage cells, this strategy bypasses the requirement for Gal4-mediated transcriptional activation and instead relies on direct cell type-specific promoter-driven expression. This avoids potential UAS-dependent transcriptional silencing and demonstrates the flexibility of the platform for use in diverse cellular and genetic contexts. As a functional validation, we targeted *slc37a2* (**Fig. 4a**), the gene mutated in the zebrafish *bubblebrain* (*blb*) mutant, in which microglia exhibit a characteristic hypertrophic morphology with enlarged phagosomes and excessive intracellular accumulation of debris (Villani et al., 2019). We reasoned that effective knockdown of *slc37a2* in wild-type microglia should recapitulate the *bubblebrain* phenotype (**Fig. 4b**).

**Fig. 4.**
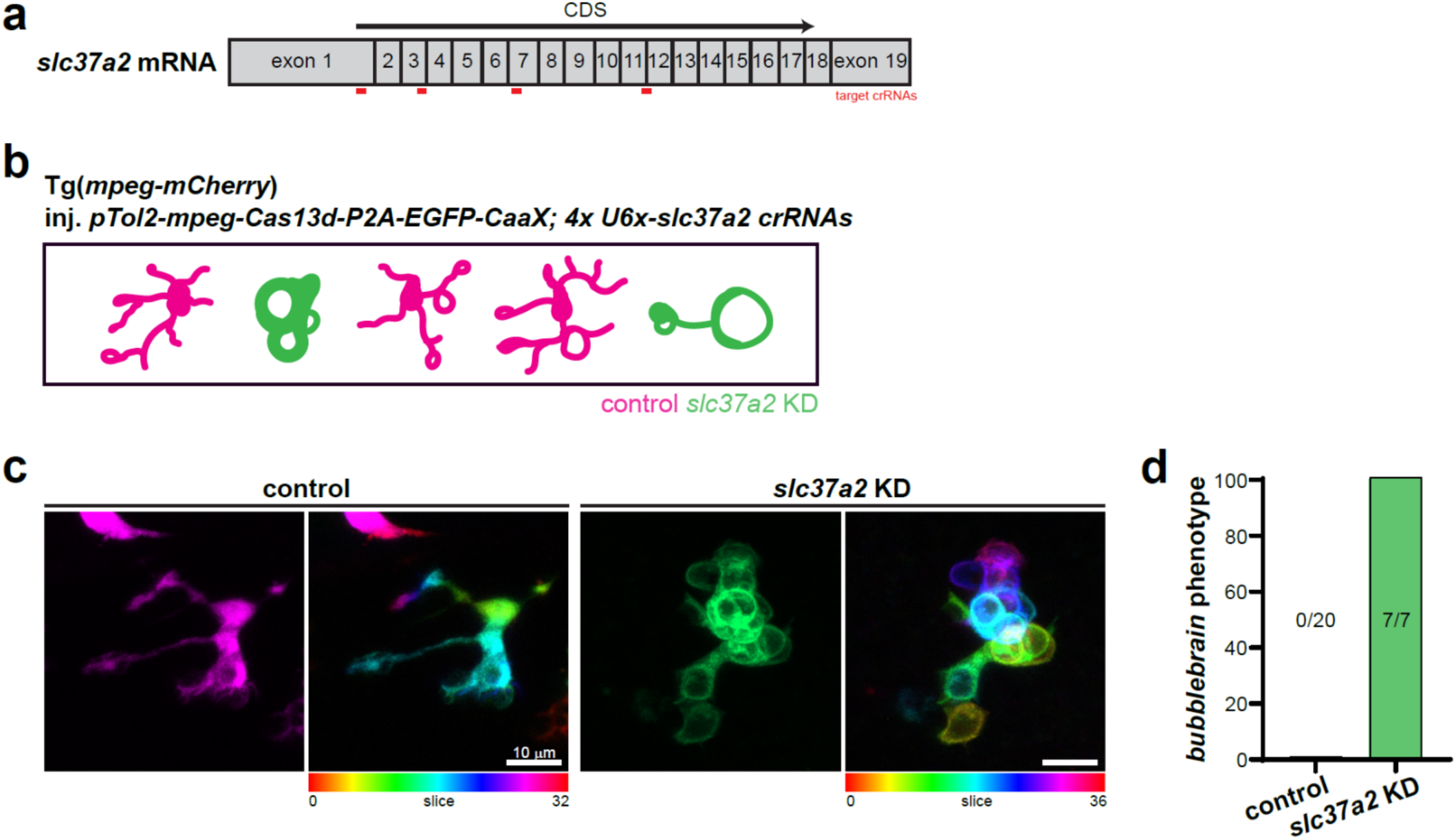
Functional validation of the cell-specific Cas13d system in microglia. **a**. Four distinct crRNAs (in red) were chosen against the protein-coding sequence (CDS) of the mature mRNA sequence of zebrafish *slc37a2* transcript. **b**. Schematic of experimental design. Single-cell embryos from Tg(*mpeg1-mCherry*) were injected with *pTol2-mpeg-Cas13d-P2A-EGFP-CaaX; 4x U6x-slc37a2 crRNAs*, which enables simultaneous visualization of mCherry+ control microglia (magenta) and EGFP+ Cas13d-targeted microglia (green) within the same animal. **c**. Representative maximum-intensity z-projected fluorescence images of control and *slc37a2* KD microglia. Pseudocolored images (right) depict individual optical z-slices in distinct colors to enhance visualization of individual phagosomes that accumulate in *slc37a2* KD microglia. **d**. Quantification of the *bubblebrain* phenotype in control and *slc37a2* KD microglia. N=20 microglia/control, N=7 microglia/*slc37a2* KD.

We injected *pTol2-mpeg1-Cas13d-P2A-EGFP-CaaX; 4x U6x-slc37a2 crRNAs* into one-cell-stage embryos of Tg(*mpeg1-mCherry*) line. This strategy enabled direct comparison of Cas13d-positive (EGFP^+^) and Cas13d-negative (mCherry^+^EGFP^-^) microglia within the same brain at 5-6 dpf. Notably, all EGFP^+^, Cas13d^-^expressing microglia with *slc37a2* knockdown displayed the *bubblebrain* phenotype, characterized by pronounced cellular enlargement and the presence of multiple expanded phagosomes (**Fig. 4c,d**). The complete penetrance of this phenotype among targeted cells indicates highly efficient and cell-autonomous gene knockdown in microglia. These results validate the efficacy of our Cas13d system in microglia and establish a rapid, cell type-specific genetic platform for interrogating microglial gene function *in vivo*. The high penetrance and single-cell resolution achieved here provide a robust foundation for targeted and multiplexed genetic approaches aimed at elucidating the molecular mechanisms governing microglial development and function within an intact vertebrate nervous system.

## Discussion

Here we present an all-in-one, cell type-specific Cas13d system for zebrafish that enables efficient and quantifiable RNA knockdown *in vivo*. By combining RfxCas13d with the cell type-specific KalTA4/UAS binary expression system and Tol2-mediated genomic integration, we established a platform that reliably depletes target transcripts and produces reproducible phenotypes in astrocytes and oligodendrocyte lineage cells. By driving Cas13d expression under the microglia/macrophage-specific *mpeg1* promoter, we further demonstrate that this system is readily adaptable to microglia, a cell population of increasing interest in both neurodevelopmental and neurodegenerative disorders that has historically been difficult to genetically target. Collectively, this platform addresses a longstanding limitation in zebrafish neurobiology: the lack of robust tools for cell type-specific loss-of-function studies with a reliable validation strategy.

While prior Cas9-based systems have enabled transformative advances in zebrafish genetics, cell type-specific gene disruption using DNA-targeting nucleases is frequently complicated by mosaic in/del formation, incomplete penetrance, and challenges in validating protein depletion, particularly in the absence of reliable antibodies. Moreover, genomic disruption can trigger transcriptional compensation or long-term adaptive responses that obscure primary gene function. In contrast, RNA-targeting with Cas13d enables direct mRNA transcript depletion without altering genomic DNA, providing a reproducible and quantifiable knockdown strategy that circumvents variability associated with genomic DNA editing. The use of stable Tol2-mediated integration further enhances consistency and enables phenotypic analyses across generations, thereby supporting both hypothesis-driven studies and high-throughput screening applications. Together, these features substantially expand the genetic toolkit available for dissecting glial cell biology *in vivo*.

The modular design of our construct provides flexibility for future refinement and broader applications. Multiplex gene targeting can be readily achieved by incorporating different U6x-driven crRNA cassettes, enabling simultaneous and combinatorial knockdown of multiple genes or distinct transcript regions. This feature is particularly advantageous in zebrafish neurobiology research, where whole-genome duplication has resulted in extensive genetic redundancy and paralog compensation. Temporal control of gene knockdown may be introduced by pairing this platform with inducible systems, such as Gal4-EcR, tTA/TRE, or heat-shock promoters, thereby facilitating the dissection of developmental stage-specific gene function. In addition, compatibility with alternative binary expression systems, including QF/QUAS, may further extend the range of cell types and experimental paradigms accessible using this approach.

The cell type-specific Cas13d platform described here is readily compatible with *in vivo* time-lapse imaging, quantitative morphological analyses, and integration with functional reporters, including calcium indicators, metabolic sensors, and transcriptional reporters. Because targeted and non-targeted cells can be analyzed within the same tissue with different fluorescent reporters, this system enables internal controls at single-cell resolution, reducing inter-animal variability and increasing statistical power. The ability to directly compare manipulated and neighboring unmanipulated cells within the same animal provides a powerful framework for dissecting cell-autonomous mechanisms *in vivo*.

Despite these advantages, there remain several limitations to the system. First, the efficiency of individual crRNAs remains difficult to thoroughly predict accurately *in silico*. While computational tools such as RNAfold can inform guide design, empirical validation is often necessary to identify highly effective crRNAs, particularly when knockdown efficiency cannot be directly assessed by fluorescence *in situ* hybridization or other quantitative measures of transcript depletion. Second, because Cas13d-mediated knockdown depends on sustained expression of the transgene, highly abundant or rapidly transcribed targets may require optimization to achieve maximal depletion. Incorporating regulatory elements that enhance transcript stability and expression efficiency, such as the woodchuck hepatitis virus post-transcriptional regulatory element (WPRE), may further improve knockdown efficacy. Third, the KalTA4/UAS system in zebrafish produces mosaic expression due to epigenetic silencing of UAS-driven transgenes; as a result, it can limit the ability to perform a saturating, population-wide targeting. To overcome this constraint, Cas13d may be directly expressed under the control of cell type-specific promoters, as we have developed for microglial experiments using the *mpeg1* promoter. Finally, although optimized Cas13d variants have been reported to exhibit minimal collateral activity, the possibility of off-target or bystander RNA degradation cannot be completely excluded and should be systematically evaluated in future applications.

Looking ahead, this all-in-one, cell type-specific Cas13d platform will be useful beyond zebrafish, as Cas13d has been shown to function efficiently in both *in vitro* and *in vivo* mammalian systems. The relatively small size of Cas13d compared to Cas9 facilitates packaging into viral vectors, enabling delivery of the effector and its guide RNAs within a single viral construct. By enabling efficient, cell type-specific RNA knockdown *in vivo*, this Cas13d system complements existing genetic approaches and provides a versatile framework for investigating cell-autonomous mechanisms underlying neural development, glial cell biology, and neurological disease across species.

## Methods

### Zebrafish husbandry and maintenance

Wild-type zebrafish (AB strain), *casper*, *nacre*, Tg(*slc1a3b-KalTA4*) (Lambert et al., 2025), Tg(*UAS-myrmScarlet*) (Li et al., 2024), Tg(*sox10-KalTA4*) (Almeida & Lyons, 2015), and Tg(*mpeg1-mCherry*) (Ellett et al., 2011b) transgenic lines were used in this study. All zebrafish experiments and procedures were performed in compliance with institutional ethical regulations for animal testing and research at Oregon Health & Science University (OHSU). Experiments were approved by the Institutional Animal Care and Use Committee (IACUC) of OHSU. Zebrafish larvae and juvenile fish were nurtured using rotifer suspension and dry food (Gemma 75 and 150, respectively). Adult fish were maintained and fed with combination of brine shrimp and dry food (Gemma 300). Zebrafish in the fish facility were maintained at 28 °C with a 14 hr-10 hr light-dark cycle.

### Plasmids

To generate *pDest-Tol2-UAS-RfxCas13d-P2A-EGFP-CaaX*, the following segments were integrated into the *pDest-Tol2-4sgRNA-Cas9* Gateway destination vector using NEBuilder Hi-Fi DNA assembly (NEB): 1) UAS (For primer: CCTGGTACCCTCGAGGTCGACGG, Rev primer: GCTCGCCATGGTGGCACTAGTGGATCCCCCGGG), 2) RfxCas13d (For primer: ATCCACTAGTGCCACCATGGCGAGCGAGGC, Rev primer: GTTCGTGGCGGAATTGCCGGACACCTTCT), and 3) P2A-EGFP-CaaX (For primer: GGCAATTCCGCCACGAACTTCTCTCTGTTAAAGCA, Rev primer: GAAGGCACAGTCAGGAGAGCACACACTTGCAG). Each segment was PCR amplified using repliQa HiFi ToughMix (Quantabio). *pDest-Tol2-mpeg1-RfxCas13d-P2A-EGFP-CaaX* was similarly constructed using NEBuilder Hi-Fi DNA assembly with the following primer sets:1) mpeg1 (For primer: TAGGTCTCTCCTGGTATGTTGGAGCACATCTGACATCTGA, Rev primer: GCCATGGTGGCTTTTGCTGTCTCCTGCACTAATGT), 2) RfxCas13d-P2A-EGFP (For primer: GGGACAGCAAAAGCCACCATGGCGAGCG, Rev primer: GAGGGTTCAGCTTAGATCTTCCTCCT), 3) CaaX-pDest vector (For primer: AGGAGGAAGATCTAAGCTGAACCCT, Rev primer: ATGTGCTCCAACATACCAGGAGAGACCTAGCGTACC).

The existing *pU6x* vectors for zebrafish CRISPR-Cas9 were modified to remove the *tracrRNA* scaffold sequences at the 3′ end while preserving *BsmBI* cloning sites for insertion of *DR30-crRNA* sequences and *BsaI* sites compatible with the Gateway destination vector using Q5 Site-Directed Mutagenesis Kit (NEB). All plasmids were designed and modified using SnapGene. All constructs generated in this study were confirmed by Oxford Nanopore whole plasmid sequencing (Plasmidsaurus) and are available upon request.

### CRISPR-Cas13d targeting sequences used in this study

*s1pr1* #1: AGCCAGTGCAAACCATGGATGA

*s1pr1* #2: CGTCATCCTGTACGCCCGCATC

*s1pr1* #3: ACGAGATGCGCCGGGCCTTCAT

*s1pr1* #4: CTCGCCCAGGGAAACCATAGTG

*tardbpa* #1: CGTGTGGCGGAGGATGAAAACGA

*tardbpa* #2: AGTTTTTTATGCAGTATGGCGAA

*tardbpb* #1: CTTCATCTGCGACCAAGATCAAG

*tardbpb* #2: ACAGGAAATTCAAAAGGGTTTGG

*slc37a2* #1: ATGAAGTCTCTAGCTCCGGGCAT

*slc37a2* #2: ACCTGGTGCGATTGGGTTCCTTT

*slc37a2* #3: TTTATCATGGGAGTCTGGAATTC

*slc37a2* #4: AGGAATTTTGGGTGGCATCGTAG

### Microinjections for mosaic labeling and generation of stable transgenic lines

Fertilized zebrafish eggs at the one-cell stage were microinjected with 1 nl of an injection solution containing 30-40 ng/µl plasmid DNA and 25 ng/µl Tol2 transposase mRNA. Injected embryos were kept at 28.5 °C in 100 mm petri dishes with embryo medium until desired developmental stages for analyses or raised up to adulthood as the F_0_ generation. To generate stable lines, F_0_ fish were then out-crossed with wild-type fish and the F_1_ progenies screened for germline transmission of the fluorescent reporters at the larval stage. Transgenic-positive F_1_ carriers were further outcrossed to subsequently establish lines with a single-copy insertion.

### Whole-mount fluorescence in situ hybridization (FISH)

Whole-mount FISH was performed by following a modified version of whole-mount immuno-coupled hybridization chain reaction (WICHCR) in zebrafish embryos and larvae (Choi et al., 2018; Ibarra-García-Padilla et al., 2021). In short, 5-7 dpf larvae were fixed in 4% PFA, dehydrated in MeOH overnight, and rehydrated in PBST (0.1% Tween-20 in 1x PBS). Larvae were then permeabilized with 100% acetone at −20 °C for 12 mins. After washing and post-fixation with 4% PFA for 20 min, larval heads and tails were clipped off for efficient penetration of the probes. The samples were incubated for two overnights at 37 °C in HCR probe mixture solution (HCR probe sets + probe hybridization buffer) followed by extensive washing and incubation in amplification mixture solution (amplifier hairpins + amplification buffer) for one overnight at room temperature. The larvae were cleared in serial glycerol solutions and mounted in Aqua Polymount for confocal imaging.

*s1pr1*-B1 probe set was designed against NM_131691.3 and *egfp*-B5 probe set was designed against the *EGFP-CaaX* sequence of the all-in-one Cas13d plasmid. All FISH reagents, including B1-Alexa Fluor 647 and B5-Alexa Fluor 546, were ordered from Molecular Instruments.

### Live imaging

For live imaging, zebrafish larvae were anesthetized with 0.16 mg/ml Tricaine in embryo medium and mounted in 0.8% low-melting agarose on a 100 mm petri dish. Once solidified, embryo medium containing Tricaine (∼0.16 mg/ml) were added to cover the agarose.

### Confocal microscopy

All imaging was performed on an upright Zeiss LSM 980 confocal with Airyscan 2 in 4Y SR mode. For FISH, a 40x/1.2 Plan-Apochromat Imm Corr objective was used, and for live imaging, a 20x/1.0-NA water objective was used. All images were taken and post-processed using Zeiss Zen software.

### Quantification and statistical analysis

Astrocyte volumes were analyzed using Imaris 10 (Bitplane) software. All statistical analyses were performed with GraphPad Prism 8 software. When comparing two groups, unpaired two-tailed Student’s t-test was used. No statistical methods were used to predetermine sample sized but our sample sizes are similar to those reported in previous publications. Data distribution was assumed to be normal, and zebrafish embryos and larvae were randomly allocated to groups. Experiments were not performed blind to the conditions of the experiments; data analyses were performed blinded to the scorer or did not require manual scoring.

## Acknowledgements

We thank T. Czopka and D. Lyons for sharing transgenic lines used in the study. Imaging work was supported by the OHSU Advanced Light Microscopy Core (ALMC, RRID: SCR_009991). Special thanks to T. Perry, A. Forbes, E. Brennan, A. Reyes, K. Hamling, and S. Akram for excellent animal care and support. This work was supported by the following National Institute of Health (NIH) grants: F32NS129591 (to D.H.), F32NS123005 (to C.L.C.), R01NS124146 (to K.R.M. and M.R.F.), R21NS138718 (to K.R.M.), and R21MH133004 (to M.R.F.).

## Author contributions

D.H., C.L.C., and J.C. conceived the CRISPR-Cas13d strategy and designed experiments with input from M.R.F. and K.R.M.. D.H. and C.L.C., wrote the manuscript with input from K.R.M., M.R.F., and J.C.. D.H. and C.L.C. performed all experiments and data analyses. J.C. and D.H. designed all-in-one, cell type-specific CRISPR-Cas13d plasmid constructs, and D.H. cloned all the plasmids used in the study. D.H. generated and isolated stable transgenic fish lines for F_2_ generation experiments.

---

**Extended Data Fig. 1.**
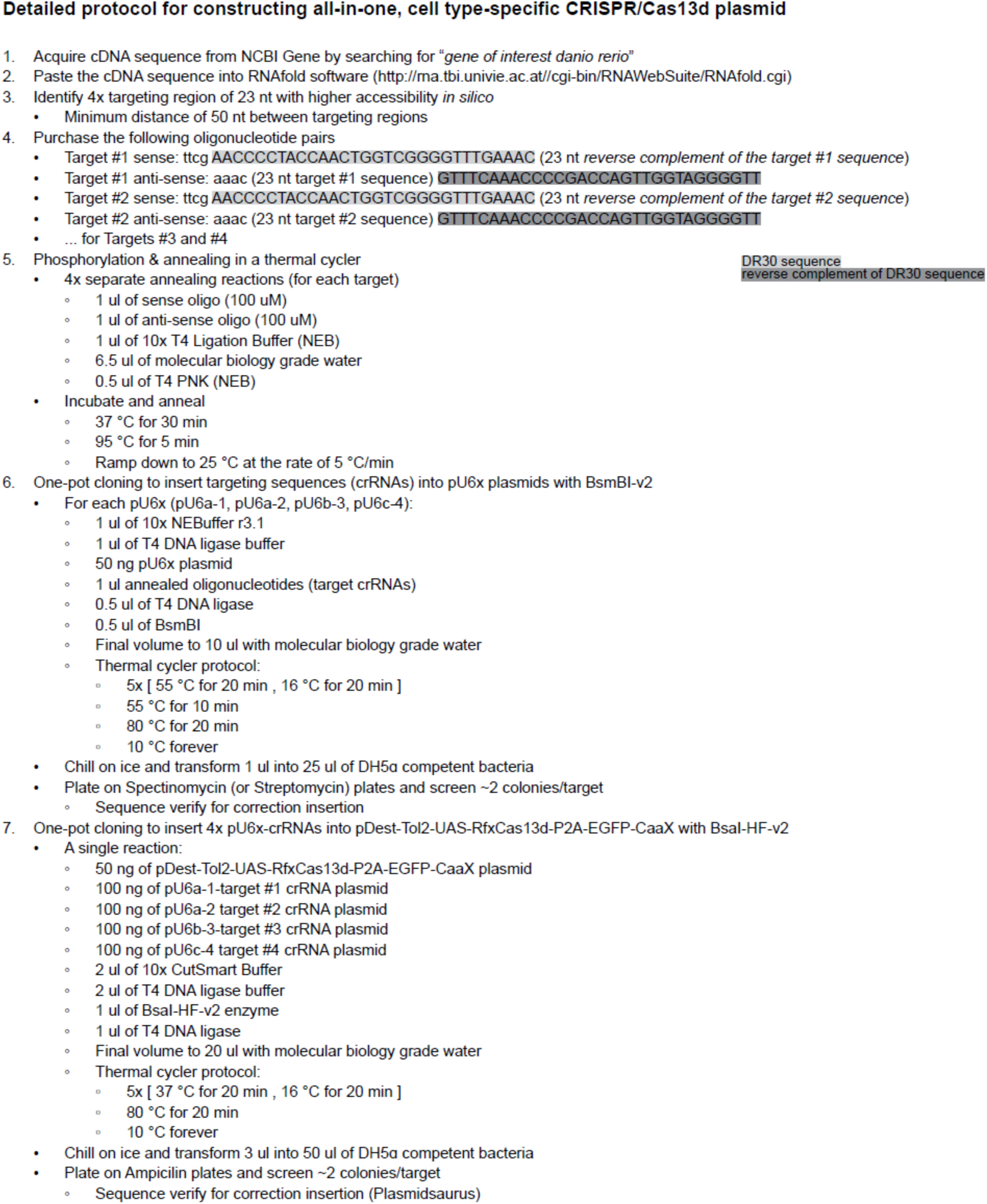
Step-by-step protocol for generating the cell type-specific Cas13d plasmid

**Extended Data Fig. 2.**
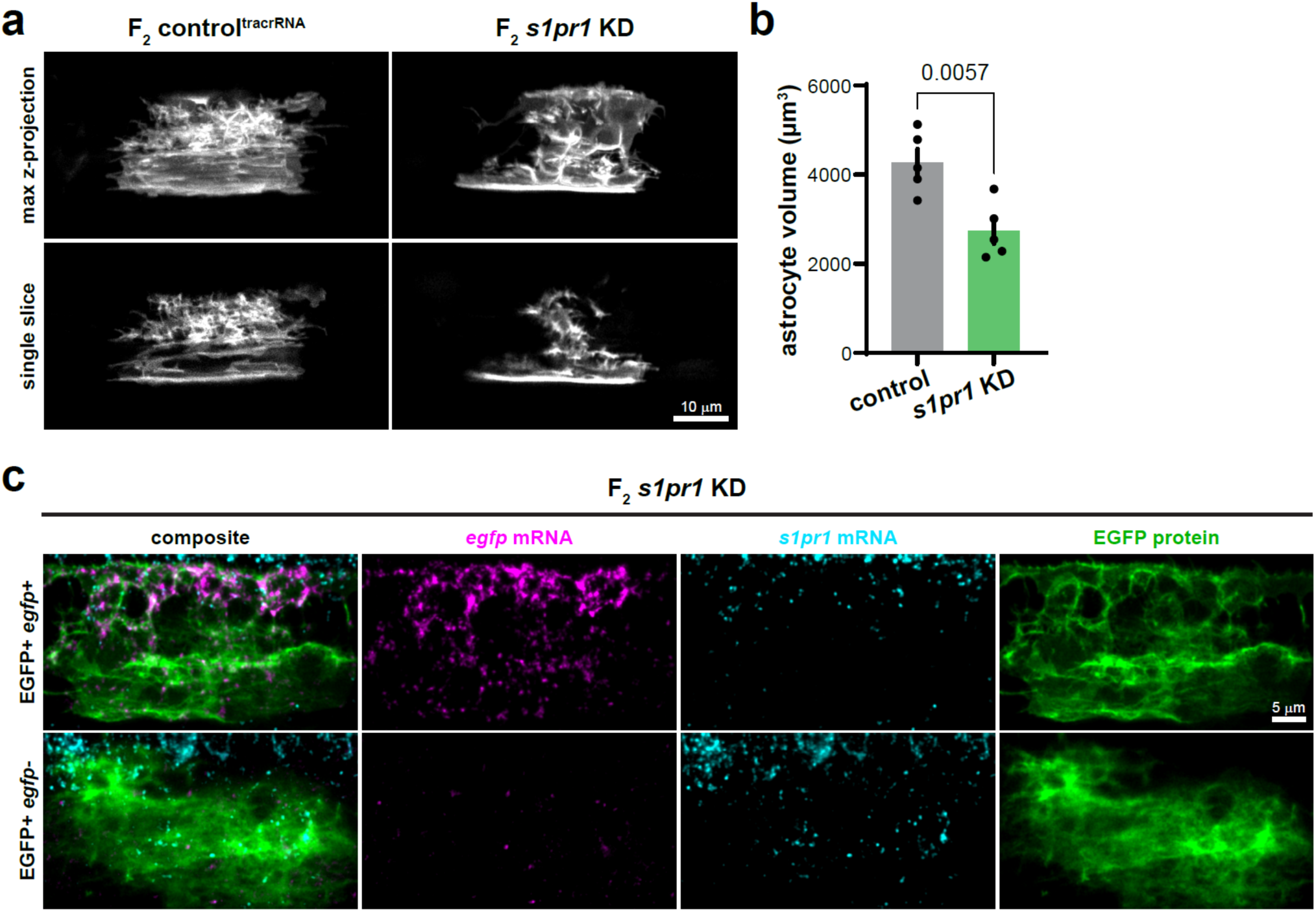
**a.** Representative images of 5-6 dpf spinal cord astrocytes in the F_2_ control^tracrRNA^ and *s1pr1* KD larval zebrafish. **b.** Quantification of astrocyte volume determined by Imaris 3D Surface rendering. N=5 astrocytes/control, N=5 astrocytes/*s1pr1* KD. **c**. Representative whole-mount FISH images of EGFP+ ventral astrocytes from F_2_ *s1pr1* KD fish.

**Extended Data Fig. 3.**
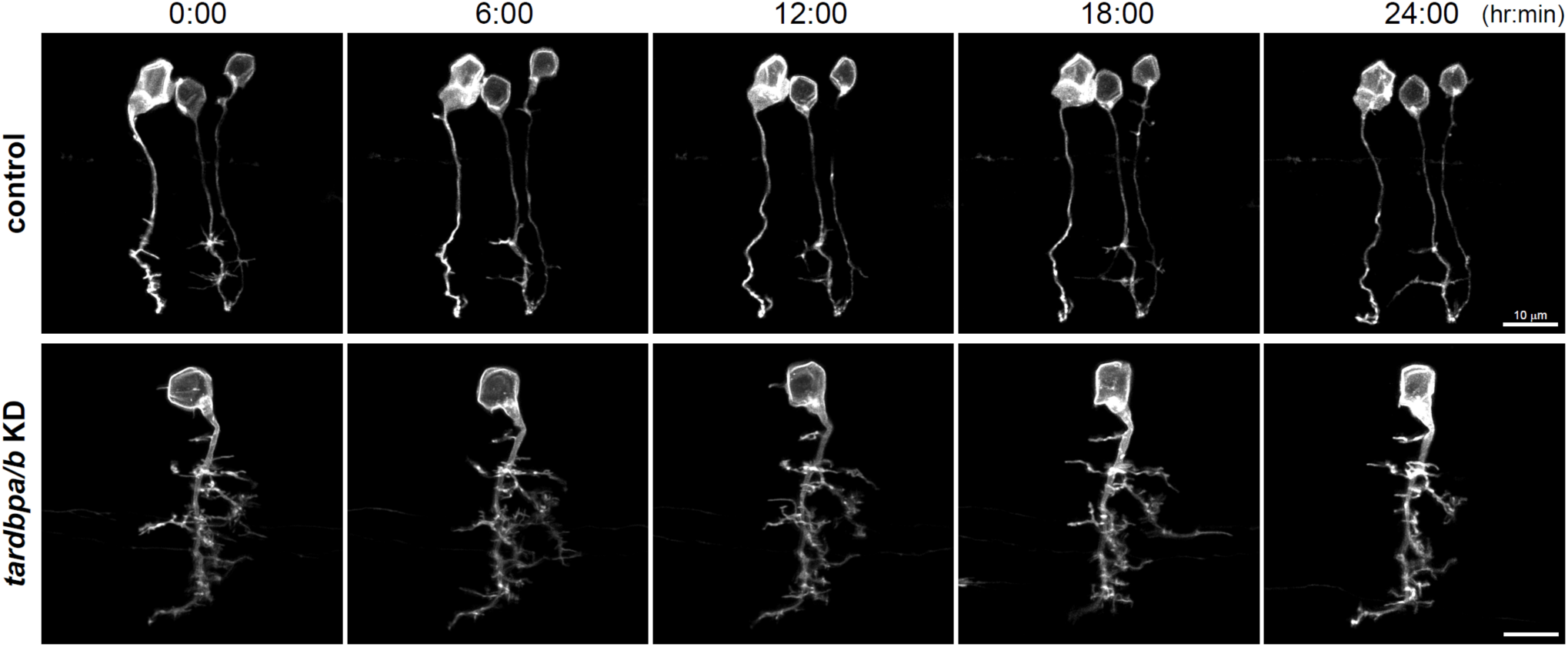
Representative images of targeted neurons at 3 dpf, imaged over 24 hours at 2-hour intervals. Neurons targeted with Tg(*sox10-KalTA4*) are resistant to Cas13d-mediated knockdown of *tardbpa/b*.

**Table 1.**
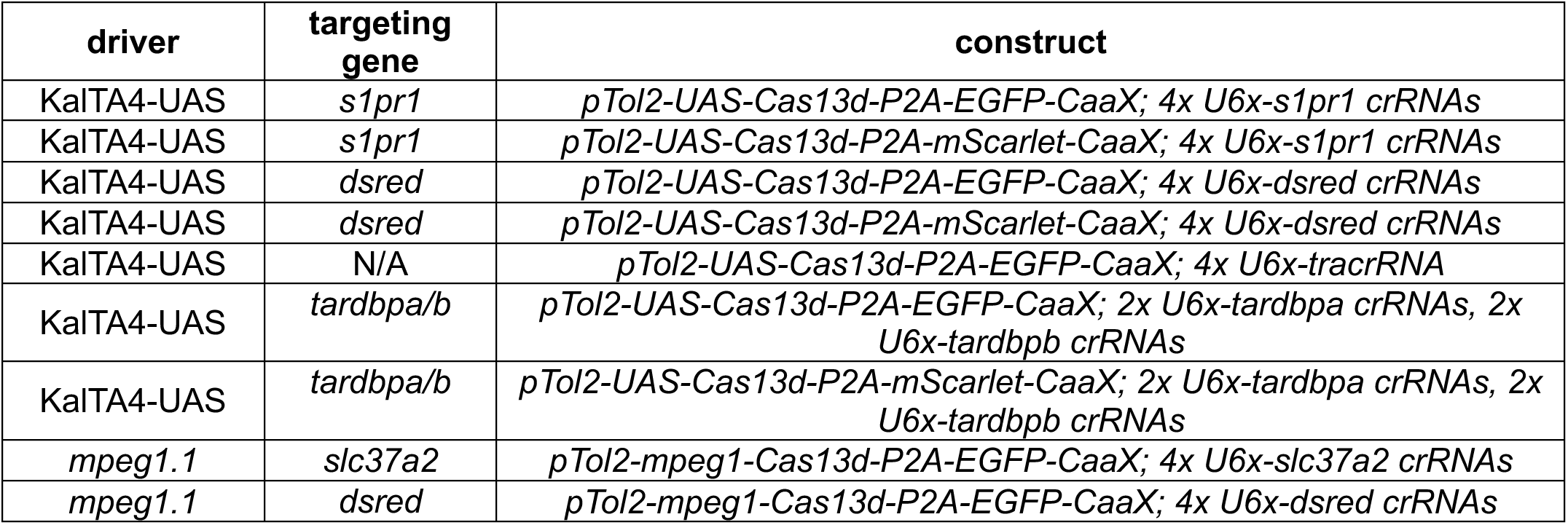
Table of available cell type-specific Cas13d expression constructs.

## References

Ablain, J., & Zon, L. I. (2016). Tissue-specific gene targeting using CRISPR/Cas9. Methods in Cell Biology, 135, 189. 10.1016/BS.MCB.2016.03.004

Abudayyeh, O. O., Gootenberg, J. S., Essletzbichler, P., Han, S., Joung, J., Belanto, J. J., Verdine, V., Cox, D. B. T., Kellner, M. J., Regev, A., Lander, E. S., Voytas, D. F., Ting, A. Y., & Zhang, F. (2017). RNA targeting with CRISPR-Cas13. Nature, 550(7675), 280–284. 10.1038/NATURE24049

Almeida, R. G., & Lyons, D. A. (2015). Intersectional Gene Expression in Zebrafish Using the Split KalTA4 System. Zebrafish, 12(6), 377–386. 10.1089/ZEB.2015.1086

Almeida, R., & Lyons, D. (2016). Oligodendrocyte Development in the Absence of Their Target Axons In Vivo. PLOS ONE, 11(10), e0164432. 10.1371/JOURNAL.PONE.0164432

Amsterdam, A., Nissen, R. M., Sun, Z., Swindell, E. C., Farrington, S., & Hopkins, N. (2004). Identification of 315 genes essential for early zebrafish development. Proceedings of the National Academy of Sciences of the United States of America, 101(35), 12792–12797. 10.1073/PNAS.0403929101

Balcaitis, S., Weinstein, J. R., Li, S., Chamberlain, J. S., & Möller, T. (2005). Lentiviral transduction of microglial cells. GLIA, 50(1), 48–55. 10.1002/GLIA.20146;PAGE:STRING:ARTICLE/CHAPTER

Bedell, V. M., Wang, Y., Campbell, J. M., Poshusta, T. L., Starker, C. G., Krug, R. G., Tan, W., Penheiter, S. G., Ma, A. C., Leung, A. Y. H., Fahrenkrug, S. C., Carlson, D. F., Voytas, D. F., Clark, K. J., Essner, J. J., & Ekker, S. C. (2012). In vivo genome editing using a high-efficiency TALEN system. Nature, 491(7422), 114–118. 10.1038/NATURE11537

Brösamle, C., & Halpern, M. E. (2002). Characterization of myelination in the developing zebrafish. Glia, 39(1), 47–57. 10.1002/GLIA.10088

Chang, N., Sun, C., Gao, L., Zhu, D., Xu, X., Zhu, X., Xiong, J. W., & Xi, J. J. (2013). Genome editing with RNA-guided Cas9 nuclease in zebrafish embryos. Cell Research, 23(4), 465–472. 10.1038/CR.2013.45

Chen, J., Poskanzer, K. E., Freeman, M. R., & Monk, K. R. (2020). Live-imaging of astrocyte morphogenesis and function in zebrafish neural circuits. Nature Neuroscience, 23(10), 1297. 10.1038/S41593-020-0703-X

Chen, J., Stork, T., Kang, Y., Nardone, K. A. M., Auer, F., Farrell, R. J., Jay, T. R., Heo, D., Sheehan, A., Paton, C., Nagel, K. I., Schoppik, D., Monk, K. R., & Freeman, M. R. (2024). Astrocyte growth is driven by the Tre1/S1pr1 phospholipid-binding G protein-coupled receptor. Neuron, 112(1), 93–112.e10. 10.1016/J.NEURON.2023.11.008/ASSET/3E7D3FAB-1ED2-442B-8674-002560C2BE63/MAIN.ASSETS/GR8.JPG

Choi, H. M. T., Schwarzkopf, M., Fornace, M. E., Acharya, A., Artavanis, G., Stegmaier, J., Cunha, A., & Pierce, N. A. (2018). Third-generation in situ hybridization chain reaction: multiplexed, quantitative, sensitive, versatile, robust. Development (Cambridge, England), 145(12). 10.1242/DEV.165753

Coombs, A. M., Heo, D., Orlin, D., Call, C. L., Bechler, M. E., Murthy, S., Emery, B., & Monk, K. R. (2026). PIEZOs regulate oligodendrocyte sheath formation, expansion, and myelination potential. BioRxiv.

Cucchiarini, M., Ren, X. L., Perides, G., & Terwilliger, E. F. (2003). Selective gene expression in brain microglia mediated via adeno-associated virus type 2 and type 5 vectors. Gene Therapy, 10(8), 657–667. 10.1038/SJ.GT.3301925

Czopka, T., ffrench-Constant, C., & Lyons, D. A. (2013). Individual Oligodendrocytes Have Only a Few Hours in which to Generate New Myelin Sheaths In Vivo. Developmental Cell, 25(6), 599–609. 10.1016/j.devcel.2013.05.013

De Santis, F., Di Donato, V., & Del Bene, F. (2016). Clonal analysis of gene loss of function and tissue-specific gene deletion in zebrafish via CRISPR/Cas9 technology. Methods in Cell Biology, 135, 171–188. 10.1016/BS.MCB.2016.03.006

Di Donato, V., De Santis, F., Auer, T. O., Testa, N., Sánchez-Iranzo, H., Mercader, N., Concordet, J. P., & Bene, F. Del. (2016). 2C-Cas9: a versatile tool for clonal analysis of gene function. Genome Research, 26(5), 681–692. 10.1101/GR.196170.115

Distel, M., Wullimann, M. F., & Köster, R. W. (2009). Optimized Gal4 genetics for permanent gene expression mapping in zebrafish. Proceedings of the National Academy of Sciences of the United States of America, 106(32), 13365–13370. 10.1073/PNAS.0903060106

Doyon, Y., McCammon, J. M., Miller, J. C., Faraji, F., Ngo, C., Katibah, G. E., Amora, R., Hocking, T. D., Zhang, L., Rebar, E. J., Gregory, P. D., Urnov, F. D., & Amacher, S. L. (2008). Heritable targeted gene disruption in zebrafish using designed zinc-finger nucleases. Nature Biotechnology, 26(6), 702–708. 10.1038/NBT1409

Driever, W., Solnica-Krezel, L., Schier, A. F., Neuhauss, S. C. F., Malicki, J., Stemple, D. L., Stainier, D. Y. R., Zwartkruis, F., Abdelilah, S., Rangini, Z., Belak, J., & Boggs, C. (1996). A genetic screen for mutations affecting embryogenesis in zebrafish. *Development (Cambridge*, England), 123, 37–46. 10.1242/DEV.123.1.37

Ellett, F., Pase, L., Hayman, J. W., Andrianopoulos, A., & Lieschke, G. J. (2011a). mpeg1 promoter transgenes direct macrophage-lineage expression in zebrafish. Blood, 117(4). 10.1182/BLOOD-2010-10-314120

Ellett, F., Pase, L., Hayman, J. W., Andrianopoulos, A., & Lieschke, G. J. (2011b). mpeg1 promoter transgenes direct macrophage-lineage expression in zebrafish. Blood, 117(4), e49–e56. 10.1182/BLOOD-2010-10-314120

Gruber, A. R., Lorenz, R., Bernhart, S. H., Neuböck, R., & Hofacker, I. L. (2008). The Vienna RNA Websuite. Nucleic Acids Research, 36(Web Server issue), W70. 10.1093/NAR/GKN188

Grunwald, D. J., & Streisinger, G. (1992). Induction of recessive lethal and specific locus mutations in the zebrafish with ethyl nitrosourea. Genetical Research, 59(2), 103–116. 10.1017/S0016672300030317

Gupta, R., Ghosh, A., Chakravarti, R., Singh, R., Ravichandiran, V., Swarnakar, S., & Ghosh, D. (2022). Cas13d: A New Molecular Scissor for Transcriptome Engineering. Frontiers in Cell and Developmental Biology, 10, 866800. 10.3389/FCELL.2022.866800

Haapaniemi, E., Botla, S., Persson, J., Schmierer, B., & Taipale, J. (2018). CRISPR–Cas9 genome editing induces a p53-mediated DNA damage response. Nature Medicine 2018 24:7, 24(7), 927–930. 10.1038/s41591-018-0049-z

Haffter, P., Granato, M., Brand, M., Mullins, M. C., Hammerschmidt, M., Kane, D. A., Odenthal, J., Van Eeden, F. J. M., Jiang, Y. J., Heisenberg, C. P., Kelsh, R. N., Furutani-Seiki, M., Vogelsang, E., Beuchle, D., Schach, U., Fabian, C., & Nüsslein-Volhard, C. (1996). The identification of genes with unique and essential functions in the development of the zebrafish, Danio rerio. *Development (Cambridge*, England), 123, 1–36. 10.1242/DEV.123.1.1

Hansen, D. V., Hanson, J. E., & Sheng, M. (2018). Microglia in Alzheimer’s disease. The Journal of Cell Biology, 217(2), 459–472. 10.1083/JCB.201709069

Hanson, K. A., Kim, S. H., & Tibbetts, R. S. (2011). RNA-Binding Proteins in Neurodegenerative Disease: TDP-43 and Beyond. Wiley Interdisciplinary Reviews. RNA, 3(2), 265. 10.1002/WRNA.111

Heo, D., Ling, J. P., Molina-Castro, G. C., Langseth, A. J., Waisman, A., Nave, K. A., Möbius, W., Wong, P. C., & Bergles, D. E. (2022). Stage-specific control of oligodendrocyte survival and morphogenesis by TDP-43. ELife, 11. 10.7554/ELIFE.75230

Herbomel, P., Thisse, B., & Thisse, C. (1999). Ontogeny and behaviour of early macrophages in the zebrafish embryo. *Development (Cambridge*, England), 126(17), 3735–3745. 10.1242/DEV.126.17.3735

Howe, K., Clark, M. D., Torroja, C. F., Torrance, J., Berthelot, C., Muffato, M., Collins, J. E., Humphray, S., McLaren, K., Matthews, L., McLaren, S., Sealy, I., Caccamo, M., Churcher, C., Scott, C., Barrett, J. C., Koch, R., Rauch, G. J., White, S., … Stemple, D. L. (2013). The zebrafish reference genome sequence and its relationship to the human genome. Nature, 496(7446), 498–503. 10.1038/NATURE12111

Hrvatin, S., Hochbaum, D. R., Nagy, M. A., Cicconet, M., Robertson, K., Cheadle, L., Zilionis, R., Ratner, A., Borges-Monroy, R., Klein, A. M., Sabatini, B. L., & Greenberg, M. E. (2018). Single-cell analysis of experience-dependent transcriptomic states in the mouse visual cortex. Nature Neuroscience, 21(1), 120–129. 10.1038/s41593-018-0112-6

Hwang, W. Y., Fu, Y., Reyon, D., Maeder, M. L., Tsai, S. Q., Sander, J. D., Peterson, R. T., Yeh, J. R. J., & Joung, J. K. (2013). Efficient genome editing in zebrafish using a CRISPR-Cas system. Nature Biotechnology, 31(3), 227–229. 10.1038/NBT.2501

Ibarra-García-Padilla, R., Howard, A. G. A., Singleton, E. W., & Uribe, R. A. (2021). A protocol for whole-mount immuno-coupled hybridization chain reaction (WICHCR) in zebrafish embryos and larvae. STAR Protocols, 2(3). 10.1016/J.XPRO.2021.100709

Jinek, M., Chylinski, K., Fonfara, I., Hauer, M., Doudna, J. A., & Charpentier, E. (2012). A programmable dual-RNA-guided DNA endonuclease in adaptive bacterial immunity. Science (New York, N.Y.), 337(6096), 816–821. 10.1126/SCIENCE.1225829

Joris, M., Schloesser, M., Baurain, D., Hanikenne, M., Muller, M., & Motte, P. (2017). Number of inadvertent RNA targets for morpholino knockdown in Danio rerio is largely underestimated: evidence from the study of Ser/Arg-rich splicing factors. Nucleic Acids Research, 45(16), 9547–9557. 10.1093/NAR/GKX638

Kirby, B. B., Takada, N., Latimer, A. J., Shin, J., Carney, T. J., Kelsh, R. N., & Appel, B. (2006). In vivo time-lapse imaging shows dynamic oligodendrocyte progenitor behavior during zebrafish development. Nature Neuroscience, 9(12), 1506–1511. 10.1038/NN1803

Kok, F. O., Shin, M., Ni, C. W., Gupta, A., Grosse, A. S., vanImpel, A., Kirchmaier, B. C., Peterson-Maduro, J., Kourkoulis, G., Male, I., DeSantis, D. F., Sheppard-Tindell, S., Ebarasi, L., Betsholtz, C., Schulte-Merker, S., Wolfe, S. A., & Lawson, N. D. (2015). Reverse genetic screening reveals poor correlation between morpholino-induced and mutant phenotypes in zebrafish. Developmental Cell, 32(1), 97–108. 10.1016/j.devcel.2014.11.018

Kosicki, M., Tomberg, K., & Bradley, A. (2018). Repair of double-strand breaks induced by CRISPR–Cas9 leads to large deletions and complex rearrangements. Nature Biotechnology 2018 36:8, 36(8), 765–771. 10.1038/nbt.4192

Kucenas, S., Snell, H., & Appel, B. (2008). nkx2.2a promotes specification and differentiation of a myelinating subset of oligodendrocyte lineage cells in zebrafish. Neuron Glia Biology, 4(2), 71–81. 10.1017/S1740925X09990123

Kushawah, G., Hernandez-Huertas, L., Abugattas-Nuñez del Prado, J., Martinez-Morales, J. R., DeVore, M. L., Hassan, H., Moreno-Sanchez, I., Tomas-Gallardo, L., Diaz-Moscoso, A., Monges, D. E., Guelfo, J. R., Theune, W. C., Brannan, E. O., Wang, W., Corbin, T. J., Moran, A. M., Sánchez Alvarado, A., Málaga-Trillo, E., Takacs, C. M., … Moreno-Mateos, M. A. (2020). CRISPR-Cas13d Induces Efficient mRNA Knockdown in Animal Embryos. Developmental Cell, 54(6), 805–817.e7. 10.1016/j.devcel.2020.07.013

Lambert, G. G., Crespo, E. L., Murphy, J., Turner, K. L., Gershowitz, E., Cunningham, M., Boassa, D., Luong, S., Celinskis, D., Allen, J. J., Venn, S., Zhu, Y., Karadas, M., Chen, J., Marisca, R., Gelnaw, H., Nguyen, D. K., Hu, J., Sprecher, B. N., … Shaner, N. C. (2025). CaBLAM: a high-contrast bioluminescent Ca2+ indicator derived from an engineered Oplophorus gracilirostris luciferase. Nature Methods 2025 23:1, 23(1), 205–215. 10.1038/s41592-025-02972-0

Li, J., Miramontes, T. G., Czopka, T., & Monk, K. R. (2024). Synaptic input and Ca2+ activity in zebrafish oligodendrocyte precursor cells contribute to myelin sheath formation. Nature Neuroscience, 27(2), 219–231. 10.1038/S41593-023-01553-8

Lv, S., Zhao, X., Ma, X., Zou, Q., Li, N., Yan, Y., Sun, L., & Song, T. (2022). Efficient and reversible Cas13d-mediated knockdown with an all-in-one lentivirus-vector. Frontiers in Bioengineering and Biotechnology, 10, 960192. 10.3389/FBIOE.2022.960192/FULL

Maes, M. E., Colombo, G., Schulz, R., & Siegert, S. (2019). Targeting microglia with lentivirus and AAV: Recent advances and remaining challenges. Neuroscience Letters, 707. 10.1016/j.neulet.2019.134310

Mehravar, M., Shirazi, A., Nazari, M., & Banan, M. (2019). Mosaicism in CRISPR/Cas9-mediated genome editing. Developmental Biology, 445(2), 156–162. 10.1016/J.YDBIO.2018.10.008

Meng, X., Noyes, M. B., Zhu, L. J., Lawson, N. D., & Wolfe, S. A. (2008). Targeted gene inactivation in zebrafish using engineered zinc-finger nucleases. Nature Biotechnology, 26(6), 695–701. 10.1038/NBT1398

Michell-Robinson, M. A., Touil, H., Healy, L. M., Owen, D. R., Durafourt, B. A., Bar-Or, A., Antel, J. P., & Moore, C. S. (2015). Roles of microglia in brain development, tissue maintenance and repair. Brain, 138(5), 1138–1159. 10.1093/brain/awv066

Miller, J. C., Tan, S., Qiao, G., Barlow, K. A., Wang, J., Xia, D. F., Meng, X., Paschon, D. E., Leung, E., Hinkley, S. J., Dulay, G. P., Hua, K. L., Ankoudinova, I., Cost, G. J., Urnov, F. D., Zhang, H. S., Holmes, M. C., Zhang, L., Gregory, P. D., & Rebar, E. J. (2011). A TALE nuclease architecture for efficient genome editing. Nature Biotechnology, 29(2), 143–150. 10.1038/NBT.1755

Moreno-Sánchez, I., Hernández-Huertas, L., Nahón-Cano, D., Martínez-García, P. M., Treichel, A. J., Gómez-Marin, C., Tomás-Gallardo, L., da Silva Pescador, G., Kushawah, G., Egidy, R., Perera, A., Díaz-Moscoso, A., Cano-Ruiz, A., Walker, J. A., Muñoz, M. J., Holden, K., Galcerán, J., Nieto, M. Á., Bazzini, A. A., & Moreno-Mateos, M. A. (2025). Enhanced RNA-targeting CRISPR-Cas technology in zebrafish. Nature Communications 2025 16:1, 16(1), 2591-. 10.1038/s41467-025-57792-9

Mullins, M. C., Hammerschmidt, M., Haffter, P., & Nüsslein-Volhard, C. (1994). Large-scale mutagenesis in the zebrafish: in search of genes controlling development in a vertebrate. Current Biology : CB, 4(3), 189–202. 10.1016/S0960-9822(00)00048-8

Nasevicius, A., & Ekker, S. C. (2000). Effective targeted gene ‘knockdown’ in zebrafish. Nature Genetics, 26(2), 216–220. 10.1038/79951

Paolicelli, R. C., Bolasco, G., Pagani, F., Maggi, L., Scianni, M., Panzanelli, P., Giustetto, M., Ferreira, T. A., Guiducci, E., Dumas, L., Ragozzino, D., & Gross, C. T. (2011). Synaptic pruning by microglia is necessary for normal brain development. Science, 333(6048), 1456–1458. 10.1126/science.1202529

Peterson, R. T., Link, B. A., Dowling, J. E., & Schreiber, S. L. (2000). Small molecule developmental screens reveal the logic and timing of vertebrate development. Proceedings of the National Academy of Sciences of the United States of America, 97(24), 12965–12969. 10.1073/PNAS.97.24.12965;WGROUP:STRING:PUBLICATION

Pogoda, H. M., Sternheim, N., Lyons, D. A., Diamond, B., Hawkins, T. A., Woods, I. G., Bhatt, D. H., Franzini-Armstrong, C., Dominguez, C., Arana, N., Jacobs, J., Nix, R., Fetcho, J. R., & Talbot, W. S. (2006). A genetic screen identifies genes essential for development of myelinated axons in zebrafish. Developmental Biology, 298(1), 118–131. 10.1016/j.ydbio.2006.06.021

Powell, J. E., Lim, C. K. W., Krishnan, R., McCallister, T. X., Saporito-Magriña, C., Zeballos, M. A., McPheron, G. D., & Gaj, T. (2022). Targeted gene silencing in the nervous system with CRISPR-Cas13. Science Advances, 8(3). 10.1126/SCIADV.ABK2485;ISSUE:ISSUE:DOI

Prinz, M., Jung, S., & Priller, J. (2019). Microglia Biology: One Century of Evolving Concepts. Cell, 179(2), 292–311. 10.1016/j.cell.2019.08.053

Sander, J. D., Cade, L., Khayter, C., Reyon, D., Peterson, R. T., Joung, J. K., & Yeh, J. R. J. (2011). Targeted gene disruption in somatic zebrafish cells using engineered TALENs. Nature Biotechnology, 29(8), 697–698. 10.1038/NBT.1934

Schafer, D. P., Lehrman, E. K., Kautzman, A. G., Koyama, R., Mardinly, A. R., Yamasaki, R., Ransohoff, R. M., Greenberg, M. E., Barres, B. A., & Stevens, B. (2012). Microglia Sculpt Postnatal Neural Circuits in an Activity and Complement-Dependent Manner. Neuron, 74(4), 691–705. 10.1016/j.neuron.2012.03.026

Schmid, B., Hruscha, A., Hogl, S., Banzhaf-Strathmann, J., Strecker, K., Van Der Zee, J., Teucke, M., Eimer, S., Hegermann, J., Kittelmann, M., Kremmer, E., Cruts, M., Solchenberger, B., Hasenkamp, L., Van Bebber, F., Van Broeckhoven, C., Edbauer, D., Lichtenthaler, S. F., & Haass, C. (2013). Loss of ALS-associated TDP-43 in zebrafish causes muscle degeneration, vascular dysfunction, and reduced motor neuron axon outgrowth. Proceedings of the National Academy of Sciences of the United States of America, 110(13), 4986–4991. 10.1073/PNAS.1218311110;WGROUP:STRING:PUBLICATION

Shah, A. N., Davey, C. F., Whitebirch, A. C., Miller, A. C., & Moens, C. B. (2015). Rapid reverse genetic screening using CRISPR in zebrafish. Nature Methods, 12(6), 535–540. 10.1038/NMETH.3360

Shembrey, C., Yang, R., Casan, J., Hu, W., Chen, H., Singh, G. J., Sadras, T., Prasad, K., Shortt, J., Johnstone, R. W., Trapani, J. A., Ekert, P. G., & Fareh, M. (2024). Principles of CRISPR-Cas13 mismatch intolerance enable selective silencing of point-mutated oncogenic RNA with single-base precision. Science Advances, 10(51), eadl0731. 10.1126/SCIADV.ADL0731

Shin, H. Y., Wang, C., Lee, H. K., Yoo, K. H., Zeng, X., Kuhns, T., Yang, C. M., Mohr, T., Liu, C., & Hennighausen, L. (2017). CRISPR/Cas9 targeting events cause complex deletions and insertions at 17 sites in the mouse genome. Nature Communications, 8. 10.1038/NCOMMS15464

Smits, A. H., Ziebell, F., Joberty, G., Zinn, N., Mueller, W. F., Clauder-Münster, S., Eberhard, D., Fälth Savitski, M., Grandi, P., Jakob, P., Michon, A. M., Sun, H., Tessmer, K., Bürckstümmer, T., Bantscheff, M., Steinmetz, L. M., Drewes, G., & Huber, W. (2019). Biological plasticity rescues target activity in CRISPR knock outs. Nature Methods 2019 16:11, 16(11), 1087–1093. 10.1038/s41592-019-0614-5

Stainier, D. Y. R., Raz, E., Lawson, N. D., Ekker, S. C., Burdine, R. D., Eisen, J. S., Ingham, P. W., Schulte-Merker, S., Yelon, D., Weinstein, B. M., Mullins, M. C., Wilson, S. W., Ramakrishnan, L., Amacher, S. L., Neuhauss, S. C. F., Meng, A., Mochizuki, N., Panula, P., & Moens, C. B. (2017). Guidelines for morpholino use in zebrafish. PLoS Genetics, 13(10). 10.1371/JOURNAL.PGEN.1007000

Sternberg, S. H., Redding, S., Jinek, M., Greene, E. C., & Doudna, J. A. (2014). DNA interrogation by the CRISPR RNA-guided endonuclease Cas9. Nature, 507(7490), 62–67. 10.1038/NATURE13011

Streisinger, G., Walker, C., Dower, N., Knauber, D., & Singer, F. (1981). Production of clones of homozygous diploid zebra fish (Brachydanio rerio). Nature, 291(5813), 293–296. 10.1038/291293a0

Tuladhar, R., Yeu, Y., Tyler Piazza, J., Tan, Z., Rene Clemenceau, J., Wu, X., Barrett, Q., Herbert, J., Mathews, D. H., Kim, J., Hyun Hwang, T., & Lum, L. (2019). CRISPR-Cas9-based mutagenesis frequently provokes on-target mRNA misregulation. Nature Communications 2019 10:1, 10(1), 4056-. 10.1038/s41467-019-12028-5

Urnov, F. D., Miller, J. C., Lee, Y. L., Beausejour, C. M., Rock, J. M., Augustus, S., Jamieson, A. C., Porteus, M. H., Gregory, P. D., & Holmes, M. C. (2005). Highly efficient endogenous human gene correction using designed zinc-finger nucleases. Nature, 435(7042), 646–651. 10.1038/NATURE03556

Villani, A., Benjaminsen, J., Moritz, C., Henke, K., Hartmann, J., Norlin, N., Richter, K., Schieber, N. L., Franke, T., Schwab, Y., & Peri, F. (2019). Clearance by Microglia Depends on Packaging of Phagosomes into a Unique Cellular Compartment. Developmental Cell, 49(1), 77–88.e7. 10.1016/j.devcel.2019.02.014

Walker, C., & Streisinger, G. (1983). Induction of Mutations by gamma-Rays in Pregonial Germ Cells of Zebrafish Embryos. Genetics, 103(1), 125–136. 10.1093/GENETICS/103.1.125

Wang, J., Ho, W. Y., Lim, K., Feng, J., Tucker-kellogg, G., Nave, K. A., Ling, S. C., Yun, W., Lim, K., Feng, J., Tucker-kellogg, G., Nave, K. A., & Ling, S. C. (2018). Cell-autonomous requirement of TDP-43, an ALS/FTD signature protein, for oligodendrocyte survival and myelination. Proceedings of the National Academy of Sciences of the United States of America, 115(46), E10941–E10950. 10.1073/pnas.1809821115

Wessels, H. H., Méndez-Mancilla, A., Guo, X., Legut, M., Daniloski, Z., & Sanjana, N. E. (2020). Massively parallel Cas13 screens reveal principles for guide RNA design. Nature Biotechnology, 38(6), 722–727. 10.1038/S41587-020-0456-9

Wessels, H. H., Stirn, A., Méndez-Mancilla, A., Kim, E. J., Hart, S. K., Knowles, D. A., & Sanjana, N. E. (2024). Prediction of on-target and off-target activity of CRISPR-Cas13d guide RNAs using deep learning. Nature Biotechnology, 42(4), 628–637. 10.1038/S41587-023-01830-8

Wienholds, E., van Eeden, F., Kosters, M., Mudde, J., Plasterk, R. H. A., & Cuppen, E. (2003). Efficient Target-Selected Mutagenesis in Zebrafish. Genome Research, 13(12), 2700. 10.1101/GR.1725103

Xu, J., Zhu, L., He, S., Wu, Y., Jin, W., Yu, T., Qu, J. Y., & Wen, Z. (2015). Temporal-Spatial Resolution Fate Mapping Reveals Distinct Origins for Embryonic and Adult Microglia in Zebrafish. Developmental Cell, 34(6), 632–641. 10.1016/j.devcel.2015.08.018

Yin, L., Maddison, L. A., Li, M., Kara, N., Lafave, M. C., Varshney, G. K., Burgess, S. M., Patton, J. G., & Chen, W. (2015). Multiplex conditional mutagenesis using transgenic expression of Cas9 and sgRNAs. Genetics, 200(2), 431–441. 10.1534/GENETICS.115.176917/-/DC1

Zhang, Y., Chen, K., Sloan, S. A., Bennett, M. L., Scholze, A. R., O’Keeffe, S., Phatnani, H. P., Guarnieri, P., Caneda, C., Ruderisch, N., Deng, S., Liddelow, S. A., Zhang, C., Daneman, R., Maniatis, T., Barres, B. A., Wu, J. Q., Paolo Guarnieri, X., Caneda, C., … Jia Qian Wu, X. (2014). An RNA-Sequencing Transcriptome and Splicing Database of Glia, Neurons, and Vascular Cells of the Cerebral Cortex. Journal of Neuroscience, 34(36), 11929–11947. 10.1523/JNEUROSCI.1860-14.2014

