## Supplementary Figures for "All-in-one, Cas13d-based cell-specific gene knockdown system for zebrafish"

**Detailed protocol for constructing all-in-one, cell type-specific CRISPR/Cas13d plasmid**

1. Acquire cDNA sequence from NCBI Gene by searching for "*gene of interest danio rerio*"
2. Paste the cDNA sequence into RNAfold software (http://rna.tbi.univie.ac.at//cgi-bin/RNAWebSuite/RNAfold.cgi)
3. Identify 4x targeting region of 23 nt with higher accessibility *in silico*
   - Minimum distance of 50 nt between targeting regions
4. Purchase the following oligonucleotide pairs
   - Target #1 sense: ttcg AACCCCTACCAACTGGTCGGGGTTTGAAAC (23 nt *reverse complement of the target #1 sequence*)
   - Target #1 anti-sense: aaac (23 nt target #1 sequence) GTTTCAAACCCCGACCAGTTGGTAGGGGTT
   - Target #2 sense: ttcg AACCCCTACCAACTGGTCGGGGTTTGAAAC (23 nt *reverse complement of the target #2 sequence*)
   - Target #2 anti-sense: aaac (23 nt target #2 sequence) GTTTCAAACCCCGACCAGTTGGTAGGGGTT
   - ... for Targets #3 and #4
5. Phosphorylation & annealing in a thermal cycler
   - 4x separate annealing reactions (for each target)
     - 1 ul of sense oligo (100 uM)
     - 1 ul of anti-sense oligo (100 uM)
     - 1 ul of 10x T4 Ligation Buffer (NEB)
     - 6.5 ul of molecular biology grade water
     - 0.5 ul of T4 PNK (NEB)
   - Incubate and anneal
     - 37 °C for 30 min
     - 95 °C for 5 min
     - Ramp down to 25 °C at the rate of 5 °C/min
6. One-pot cloning to insert targeting sequences (crRNAs) into pU6x plasmids with BsmBI-v2
   - For each pU6x (pU6a-1, pU6a-2, pU6b-3, pU6c-4):
     - 1 ul of 10x NEBuffer r3.1
     - 1 ul of T4 DNA ligase buffer
     - 50 ng pU6x plasmid
     - 1 ul annealed oligonucleotides (target crRNAs)
     - 0.5 ul of T4 DNA ligase
     - 0.5 ul of BsmBI
     - Final volume to 10 ul with molecular biology grade water
     - Thermal cycler protocol:
       - 5x [ 55 °C for 20 min , 16 °C for 20 min ]
       - 55 °C for 10 min
       - 80 °C for 20 min
       - 10 °C forever
   - Chill on ice and transform 1 ul into 25 ul of DH5α competent bacteria
   - Plate on Spectinomycin (or Streptomycin) plates and screen ~2 colonies/target
     - Sequence verify for correction insertion
7. One-pot cloning to insert 4x pU6x-crRNAs into pDest-Tol2-UAS-RfxCas13d-P2A-EGFP-CaaX with BsaI-HF-v2
   - A single reaction:
     - 50 ng of pDest-Tol2-UAS-RfxCas13d-P2A-EGFP-CaaX plasmid
     - 100 ng of pU6a-1-target #1 crRNA plasmid
     - 100 ng of pU6a-2 target #2 crRNA plasmid
     - 100 ng of pU6b-3-target #3 crRNA plasmid
     - 100 ng of pU6c-4 target #4 crRNA plasmid
     - 2 ul of 10x CutSmart Buffer
     - 2 ul of T4 DNA ligase buffer
     - 1 ul of BsaI-HF-v2 enzyme
     - 1 ul of T4 DNA ligase
     - Final volume to 20 ul with molecular biology grade water
     - Thermal cycler protocol:
       - 5x [ 37 °C for 20 min , 16 °C for 20 min ]
       - 80 °C for 20 min
       - 10 °C forever
   - Chill on ice and transform 3 ul into 50 ul of DH5α competent bacteria
   - Plate on Ampicilin plates and screen ~2 colonies/target
     - Sequence verify for correction insertion (Plasmidsaurus)

DR30 sequence
reverse complement of DR30 sequence

**Extended Data Fig. 1**
Step-by-step protocol for generating the cell type-specific Cas13d plasmid

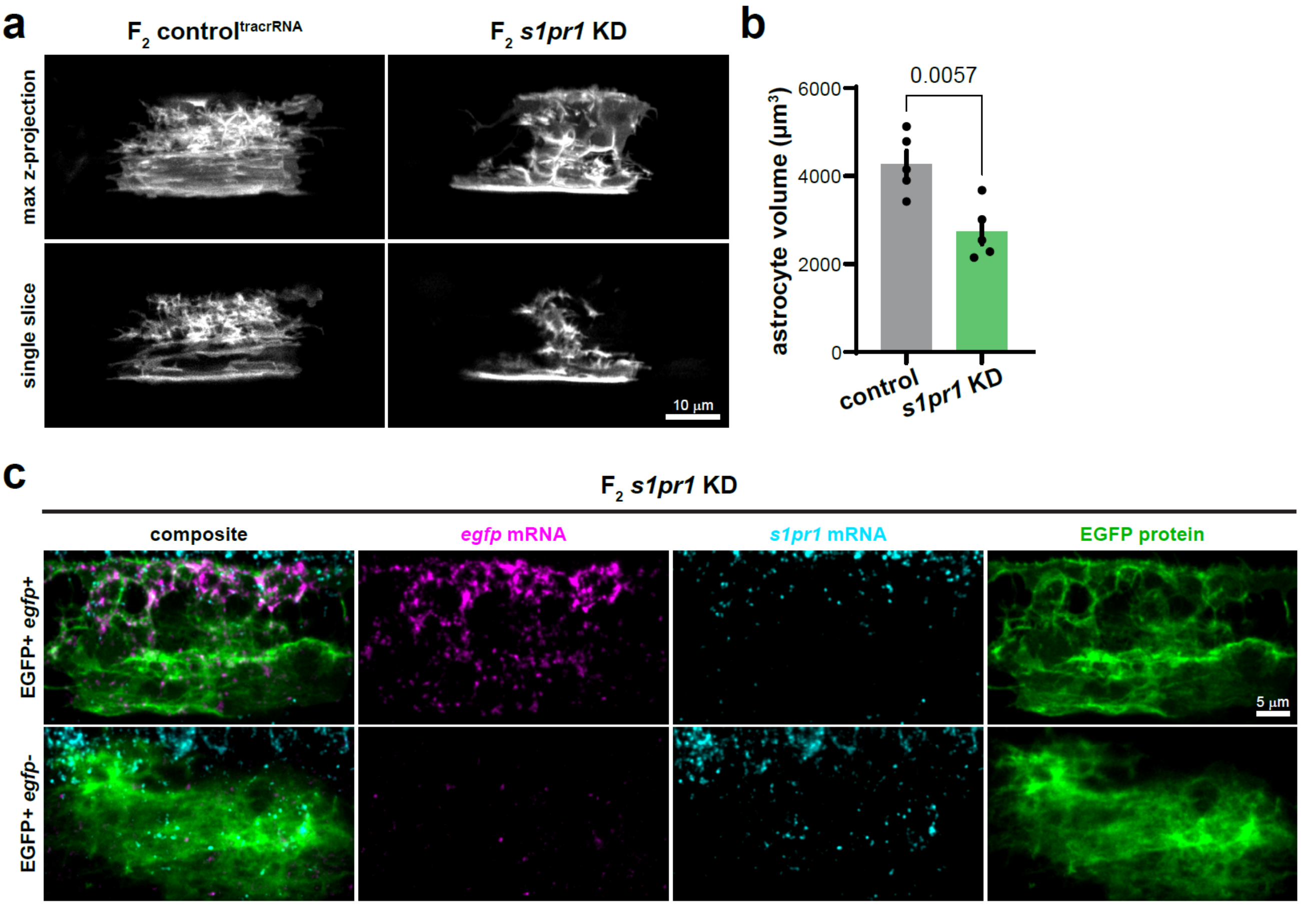


**Extended Data Fig. 2**

**a**. Representative images of 5-6 dpf spinal cord astrocytes in the $F_2$ control$^{tracrRNA}$ and *s1pr1* KD larval zebrafish. **b**. Quantification of astrocyte volume determined by Imaris 3D Surface rendering. N=5 astrocytes/control, N=5 astrocytes/*s1pr1* KD. **c**. Representative whole-mount FISH images of EGFP+ ventral astrocytes from $F_2$ *s1pr1* KD fish.

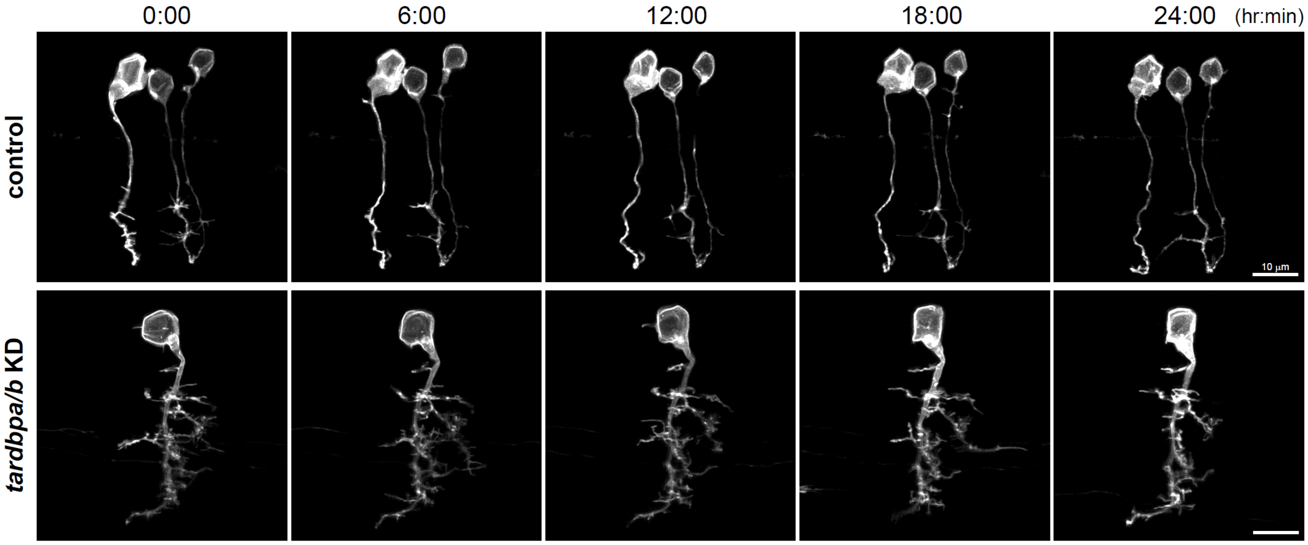


**Extended Data Fig. 3**
Representative images of targeted neurons at 3 dpf, imaged over 24 hours at 2-hour intervals. Neurons targeted with Tg(*sox10-KalTA4*) are resistant to Cas13d-mediated knockdown of *tardbpa/b*.

| driver | targeting gene | construct |
|---|---|---|
| KalTA4-UAS | *s1pr1* | *pTol2-UAS-Cas13d-P2A-EGFP-CaaX; 4x U6x-s1pr1 crRNAs* |
| KalTA4-UAS | *s1pr1* | *pTol2-UAS-Cas13d-P2A-mScarlet-CaaX; 4x U6x-s1pr1 crRNAs* |
| KalTA4-UAS | *dsred* | *pTol2-UAS-Cas13d-P2A-EGFP-CaaX; 4x U6x-dsred crRNAs* |
| KalTA4-UAS | *dsred* | *pTol2-UAS-Cas13d-P2A-mScarlet-CaaX; 4x U6x-dsred crRNAs* |
| KalTA4-UAS | N/A | *pTol2-UAS-Cas13d-P2A-EGFP-CaaX; 4x U6x-tracrRNA* |
| KalTA4-UAS | *tardbpa/b* | *pTol2-UAS-Cas13d-P2A-EGFP-CaaX; 2x U6x-tardbpa crRNAs, 2x U6x-tardbpb crRNAs* |
| KalTA4-UAS | *tardbpa/b* | *pTol2-UAS-Cas13d-P2A-mScarlet-CaaX; 2x U6x-tardbpa crRNAs, 2x U6x-tardbpb crRNAs* |
| *mpeg1.1* | *slc37a2* | *pTol2-mpeg1-Cas13d-P2A-EGFP-CaaX; 4x U6x-slc37a2 crRNAs* |
| *mpeg1.1* | *dsred* | *pTol2-mpeg1-Cas13d-P2A-EGFP-CaaX; 4x U6x-dsred crRNAs* |

**Table 1. Table of available cell type-specific Cas13d expression constructs**
